# Quantitative Characterization of Budding Yeast Polarization at the Mesoscale

**DOI:** 10.64898/2026.02.06.704348

**Authors:** Marieke M. Glazenburg, Esther Geurken, Liedewij Laan

**Affiliations:** Department of Bionanoscience, Delft University of Technology, Delft, The Netherlands

## Abstract

Cellular symmetry breaking or polarization is a key process that underlies many types of cellular behavior like division, migration and differentiation. In budding yeast, polarization is required for bud site selection. Polarization of key regulator Cdc42 has been molecularly dissected for decades, yet systematic quantification of protein localization dynamics during symmetry breaking has been lacking. Here, we present a phenomenological description of polarization in terms of physical characteristics using live cell microscopy and quantitative analysis. We identify multiple stages of Cdc42 polarization and derive mesoscale observables that describe each stage, such as establishment timescales and spot morphology indicators. We then apply this description to two case studies in which the wildtype polarity network is perturbed: a small perturbation after eliminating spatial cue communication and a larger perturbation involving removal of near-essential scaffold Bem1. This confirms the applicability of the framework in non-wildtype contexts while systematically revealing differences between genetic backgrounds. Finally, we study the dynamics of other polarizing components by extending our analysis to the Gic2 PBD domain, a Cdc42-GTP biosensor, and to polarisome component Spa2, highlighting variation in spot dynamics between protein species while also showing similarities to Cdc42. This approach enables standardized comparison between diverse molecular backgrounds at the mesoscale and helps constrain future modeling efforts.

## I. INTRODUCTION

Cells have to continuously maintain their internal organization to stay alive. An example of such intracellular ordering is polarization, a symmetry breaking process through which certain cellular components organize themselves anisotropically inside the cell, such that a preferential axis is established. Polarization can be found in a wide range of biological systems, such as stem cell differentiation, embryonic development and asymmetric cell types like epithelia [1–3]. Although manifestations of cell polarity are diverse, certain underlying mechanisms and protein complexes are conserved across systems and the tree of life [4–6]. A particular type of reversible cell polarization occurs during asymmetric cell division, for which the budding yeast *Saccharomyces cerevisiae* is a prime model system.

*S. cerevisiae* divides by growing a daughter cell from the membrane of the mother cell, a process referred to as budding [7]. In order to select the unique membrane site where the new bud will form, signaling proteins need to localize into a polarized spot at the correct time during the cell cycle. These signaling proteins in turn enable the localization of downstream components, which eventually induce bud growth by directed membrane deposition through a ring-shaped septin collar [8–10]. A central regulator of polarization in budding yeast is Cdc42, a small single domain GTPase that cycles on and off the membrane by switching between a GTP-bound and a GDP-bound state [11, 12]. Recruitment of downstream polarity proteins is governed by the GTP-bound form, which is therefore often referred to as the active state. In *S. cerevisiae*, Cdc42 is essential for establishing cell polarity and, consequently, cell proliferation [13]. Since polarization is an essential step of the budding yeast life cycle, the process is precisely regulated to ensure correct timing and uniqueness of the budding site [14, 15].

Over the years, many advances have been made in the mechanistic understanding of cell polarization. This molecular understanding has been accelerated by various types of mathematical modeling. Most notably, polarization is often described using Turing type reaction-diffusion systems, similar to typical modeling of Min waves in *Escherichia coli* [11, 16–18]. Moreover, there has been increasing attention to the role of phase separation and multivalency in polarization, leading to thermodynamic models of symmetry breaking based on soft matter physics [19–21]. However, although theoretical descriptions are becoming more advanced, many different plausible reaction networks or physical mechanisms are in principle capable of symmetry breaking in some area of their parameter space [22, 23]. Current data provide little constraints to distinguish between the resulting patterns in a quantitative manner.

Strikingly, the observation that many different molecular architectures can explain polarization is supported by insights from evolution. Comparative studies have shown that the protein composition of the polarity network is highly diverse and can change significantly even between closely related species [24, 25]. Moreover, in many cases, cells can evolutionarily recover from the loss of genes that are deemed essential or nearly essential, including polarity-related genes [26, 27]. This implies that while the macroscopic cell-level phenotype of asymmetric division is robustly maintained across evolution, the microscopic composition of the cell polarization machinery can be highly variable [28]. However, it is unclear how this microscopic variability translates to the phenotype at the intermediate or mesoscale level. Specifically, the effect of molecular variation on protein localization dynamics is largely unknown, since methods to systematically compare polarization phenotypes are currently lacking.

We address this problem by generating a phenomenological description of polarization and applying this description to a variety of molecular contexts. We demonstrate a general framework to systematically obtain physical characteristics of the dynamics of polarization using live cell microscopy and quantitative analysis and use this method to map out diversity and constraints in different genetic backgrounds. We first obtain a set of observables that describe polarization of Cdc42 in wildtype cells, revealing a process of establishment, peak localization and subsequent maintenance before bud emergence. As case studies for our framework, we apply a mild perturbation to polarization by removing communication with internal asymmetries, leading to spontaneous symmetry breaking without significant growth defects. We also study the effects of a large perturbation by removing scaffold-mediated positive feedback in Cdc42 recruitment, which significantly affects cellular health and results in a variety of different phenotypes, of which premature depolarization is most dominant. Finally, we compare the dynamics of Cdc42 to two other polarizing protein species: Gic2PBD, an isolated protein domain used as a biosensor for GTP-bound Cdc42, and Spa2, an effector of Cdc42 that is part of the polarisome. This demonstrates applicability of the framework to other polarizing components, but also highlights differences in dynamics. Together, we use these results to reveal patterns of observables that tolerate variability as well as more strongly constrained characteristics at the mesoscale, paving the way for wider application to other genetic backgrounds.

## II. RESULTS

### A. Characterization of wildtype Cdc42 polarization

Image acquisition and analysis methods are described in detail in the Methods section and Supplemental Material [29]. 3D live cell timelapse microscopy is performed on cells endogenously expressing Cdc42 labeled with sfGFP in a sandwich construct (Cdc42-sfGFP^sw^ [30]) under its native promoter. Individual cells are segmented from the field of view using a neural network combined with a tracking algorithm [31]. Polarization events are manually annotated in the time series from the unpolarized state following cytokinesis until the onset of bud emergence, i.e. the first visibility of the bud. The signal in each cell is normalized with respect to its unpolarized cytoplasmic content and a signal threshold is calculated based on the variance in unpolarized cytoplasmic intensity. Using this threshold, polarized signal is differentiated from the background, yielding a 3D reconstruction of the polarized zone developing over time (Fig. 1(a)). The summed intensity inside the polar zone of a representative polarization event is shown in Fig. 1(b), as well as an effective smoothed trace obtained by applying a Savitzky-Golay filter. Combining the traces of all individual events by aligning the intensity at the onset of signal increase and taking the mean for each time point yields the average polarization curve shown in Fig. 1(c). Since not all traces are of the same length, the mean trace is cut off after the average total duration of polarization (i.e. the average time between polarity onset and bud emergence; details can be found in the Supplemental Material [29]).

**FIG. 1.**
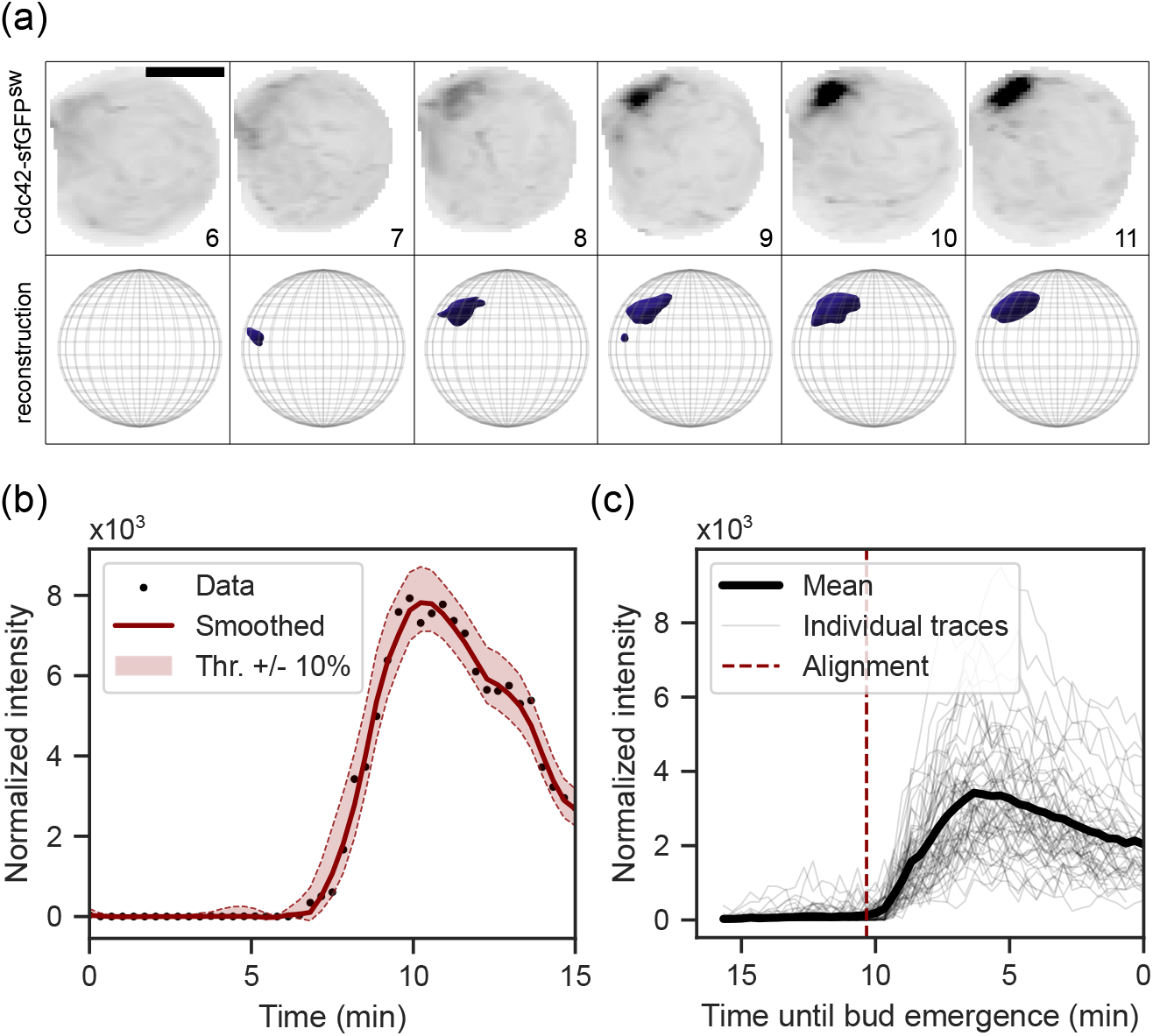
Illustration of the quantification of Cdc42 polarization in wildtype. (a) Montage of a representative polarizing wildtype cell, segmented from the background. Top row shows maximum z-projections of fluorescence imaging data of Cdc42-sfGFP^sw^, bottom row shows the corresponding 3D reconstruction after signal thresholding. Numbers indicate time in minutes relative to the start of the annotated event. Frames are snapshots; the actual time resolution is 20 s. Scale bar is 3 µm. (b) Summed normalized signal intensity inside the thresholded area of the cell from (a), with a Savitzky-Golay smoothing filter applied. Shaded area illustrates the effect of raising/lowering the signal threshold by 10%. (c) All individual unfiltered intensity traces and mean intensity trace of Cdc42 in wildtype cells (n = 61), aligned at the onset of signal increase, cut off after the average total duration of polarization.

We start by describing the general features of the thresholded polarization signal along this trajectory. As a baseline, we consider polarization of Cdc42 in wildtype cells. Examining the mean intensity profile (Fig. 2(a)), three segments can be distinguished: an establishment phase in which cellular symmetry is broken and the signal gradually increases, a peak moment when the signal reaches its maximum and a post-establishment or maintenance phase in which a slight decrease of signal is observed. We define a number of quantitative observables for each segment to characterize the properties of polarization during that stage (see Table I and Fig. 2(b)). Details behind the process of deriving each observable from the data are described in the Supplemental Material [29]. Resulting distributions for each observable are shown in Fig. 2(c).

**FIG. 2.**
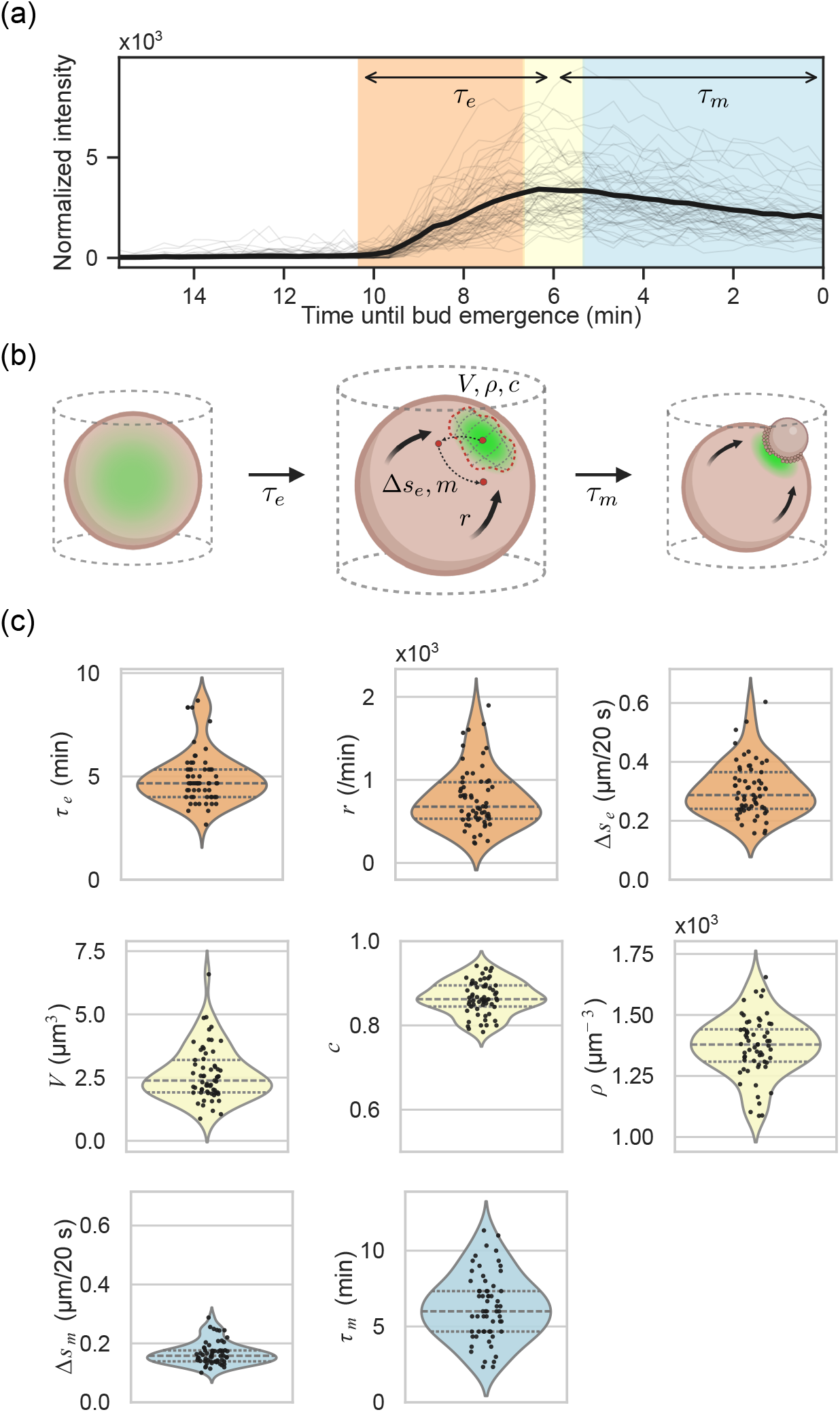
Characteristics of Cdc42 polarization in wildtype. (a) Individual normalized intensity traces (thin lines) and mean intensity (thick line) of polarized Cdc42-sfGFP^sw^ in wildtype cells (n = 61) as in figure 1d. Shading indicates stages of polarization: establishment (red), peak (yellow) and maintenance (blue). (b) Cartoon illustrating polarization of Cdc42 in budding yeast. Symbols indicate quantitative observables as defined in Table I and plotted in (c). (c) Distributions of polarization observables from Table I for Cdc42 in wildtype cells. Colors correspond to the polarization stages in (a). Dashed lines indicate data quartiles.

**Table I.** Observable definitions. Name, symbol and definition of each of the observables used to characterize polarization during the phases identified in Fig. 2(a): establishment (est.), peak and maintenance (maint.). Details for the derivations are given in the Supplemental Material [29].

|  | Observable | Symbol | Definition |
| --- | --- | --- | --- |
| *Est.* | Establishment time | $\tau_e$ | Time between onset of signal increase and signal peak |
| | Signal increase rate | $r$ | Peak total intensity divided by $\tau_e$ |
| | Mobility during establishment | $\Delta s_e$ | Average step size of the signal center of mass per 20 second time frame |
| *Peak* | Peak spot volume | $V$ | Total thresholded voxel volume |
| | Peak spot density | $\rho$ | Total intensity divided by $V$ |
| | Peak spot circularity | $c$ | Circularity of 2D projected spot contour ($0 < c \leq 1$) |
| *Maint.* | Mobility during maintenance | $\Delta s_m$ | Average step size of the signal center of mass per 20 second time frame |
| | Time before bud emergence | $\tau_m$ | Time between signal peak and bud emergence |

Together, these observables describe polarization as follows. During establishment time *τ*_*e*_, the total amount of localized Cdc42 increases at an average rate *r* until it reaches a peak. While establishment takes place, the spot has some collective mobility Δ*s*_*e*_ with which it moves along the membrane. At the peak, the spot reaches a volume *V* with signal density *ρ* and projected circularity *c*. After reaching the peak, a polarity maintenance stage begins in which the localized signal steadily decreases over a period *τ*_*m*_ until the daughter cell emerges. During this period, the spot retains some collective mobility Δ*s*_*m*_.

Several observations can be made at this stage. Firstly, the temporal dynamics of polarized Cdc42 show that apart from the initial phase of rapid recruitment, another phase characterized by slight depolarization occurs before bud emergence is visible. Where previous studies reported continuous oscillations of multiple wave periods for certain polarity proteins [32, 33], Cdc42 does not seem to follow this behavior. However, *τ*_*m*_ is typically equal to or larger than *τ*_*e*_, which together with the significant decrease in polarized signal does suggest an important contribution of negative feedback in the regulation of Cdc42 during the later stages of polarization. Secondly, the spot mobility Δ*s*_*m*_ during maintenance is roughly two-fold lower than the initial mobility Δ*s*_*e*_, indicating stabilization of Cdc42 positioning at this stage. This is consistent with earlier results where GTP-bound Cdc42 was observed to initially drift along the membrane, while mobility sharply decreased later in polarization [34].

It is worth noting that, although the average behavior described here is highly reproducible, there is significant heterogeneity between individual cells in the population. This is evident by the large spreads in the distributions in Fig. 2(c). Hence, although polarization is an essential and strongly conserved process, the physical characteristics of polarity establishment can vary significantly even between healthy cells. This suggests that polarization tolerates a degree of phenotypic variability, despite strong biological constraints on timing and site uniqueness.

### B. Small perturbation: dynamics of spontaneous symmetry breaking

After obtaining a quantitative description of Cdc42 polarization in a wildtype background, we set out to apply this framework to a case in which polarization is lightly perturbed, with little effect on cellular health. We opted for the relatively well-studied case of spontaneous symmetry breaking, which occurs when communication with internal spatial landmarks is removed. In wildtype budding yeast, localization of Cdc42 is primed by the presence of internal landmark complexes that are formed as a consequence of the previous division event: the bud scar in the mother and the birth scar in the daughter cell [7, 8]. In haploid cells, the next polarization event will occur along this ring-shaped landmark located at the proximal pole of the cell. Diploid cells can additionally polarize along a second landmark complex located directly opposite to the bud scar at the distal pole [35]. However, in absence of these spatial cues, cells are still able to polarize and thus spontaneously break their symmetry with no significant effects on overall growth rate of the population [36–39]. Removing communication with spatial cues in haploid cells can be achieved by knocking out the GTPase gene *RSR1*, which interacts with both the bud scar complex and the polarization machinery (Fig. 3(a)) [34, 40–43]. However, previous work revealed undesired side effects of *RSR1* deletion and suggested an alternative approach by knocking out *AXL2* and *RAX1* [44], which are direct components of the proximal and distal landmark complexes respectively. For our purposes, we opted to include both mutant variants.

**FIG. 3.**
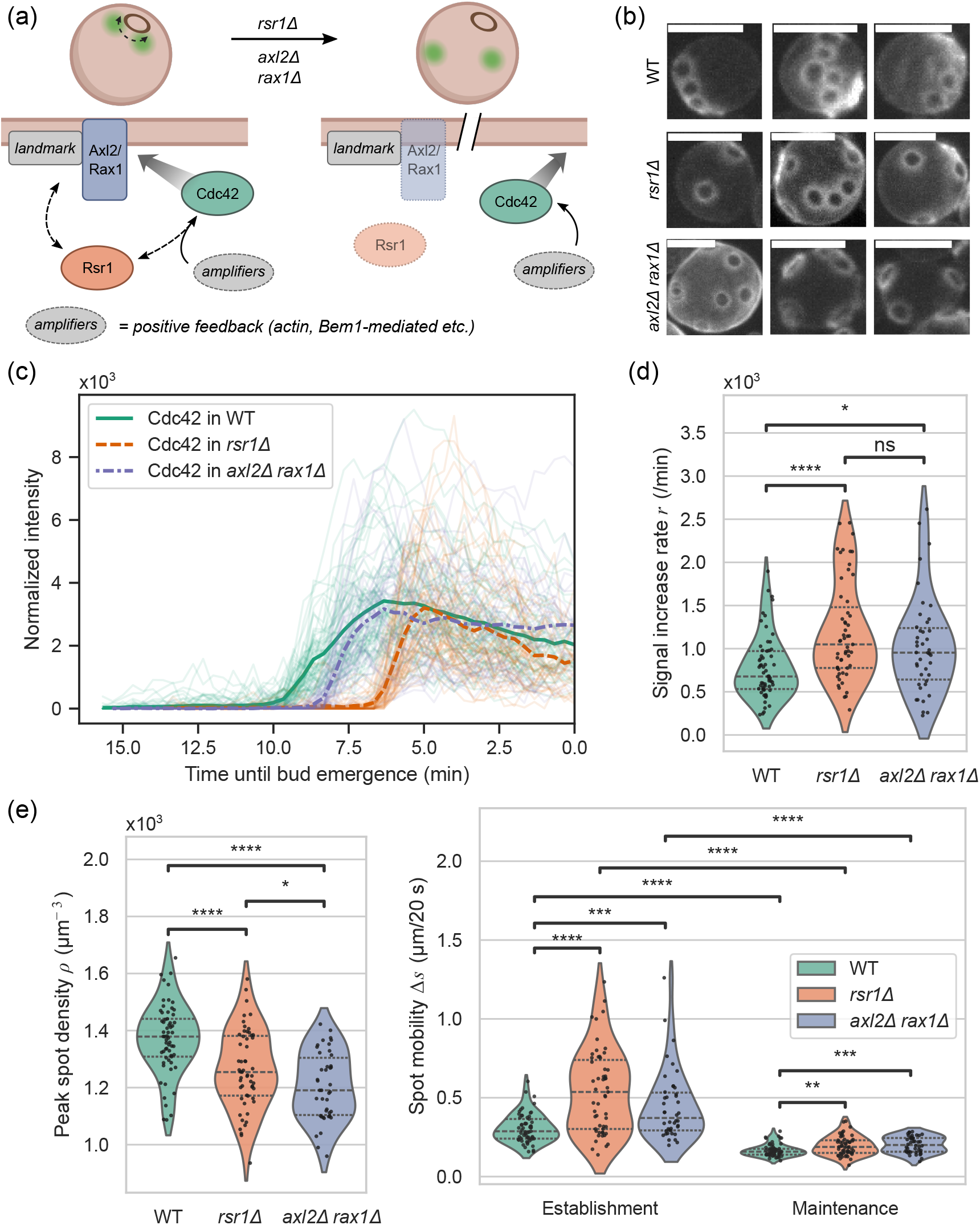
Characteristics of Cdc42 polarization in spatial cue mutants. (a) Cartoon illustrating the role of Rsr1, Axl2 and Rax1 in polarization. Spot site selection in wildtype cells is linked to internal landmark complexes, which contain Axl2 and Rax1, through the GTPase Rsr1, which then interacts with positive feedback inducing components (indicated as *amplifiers*) that stimulate further localization of Cdc42 to the polar zone. Removing either both landmark components Axl2 and Rax1 or the communication link Rsr1 removes the bias towards the landmark, allowing amplification of Cdc42 anywhere on the membrane. (b) Z-projected images of representative cells stained with calcofluor white. Scale bars 5 µm. (c) Individual normalized intensity traces of polarized Cdc42-sfGFP^sw^ (thin lines) and mean intensity for wildtype (n = 61), *rsr1* Δ (n = 55) and *axl2* Δ *rax1* Δ (n = 42) cells, aligned at the end of the time series corresponding to bud emergence. (d) Distributions of signal increase rate *r* for wildtype and mutants. (e) Distributions of peak spot density *ρ* for wildtype and mutants. (f) Distributions of spot mobility during establishment Δ*s*_*e*_ and during maintenance Δ*s*_*m*_ for wildtype and mutants. All plots: \**p* < 0.05; \*\**p* < 0.01; \*\*\**p* < 0.001; \*\*\*\**p* < 0.0001; ns, not significant (Mann-Whitney U-test). Dashed lines indicate data quartiles.

The increased randomness of the budding pattern in both mutants is illustrated in Fig. 3(b). Each panel shows staining of wildtype or mutant cells with calcofluor white to visualize the cell walls and bud scar complexes on the outside of the cell resulting from previous division events. Bud scars in wildtype are chained together, indicating that the new bud site forms next to the old division site. In the mutants however, the distribution of bud scars is scattered across the cortex, showing how the new bud sites can be chosen independently from bud scar landmarks. In the following, we will characterize polarization of Cdc42 for both spatial cue mutants, using the same endogenously expressed Cdc42-sfGFP^sw^ construct as in the wildtype background.

Mean intensity traces of polarized Cdc42 in the *rsr1*Δ and the *axl2*Δ*rax1*Δ mutants, as well as a wildtype reference, are shown in Fig. 3(c). Individual intensity traces per background are aligned at the onset of polarity, as described in section II A; the mean traces are then plotted to align at the end of the timelapse corresponding to bud emergence. From this, we observe that the time between the onset of Cdc42 polarization and bud emergence (*τ*_*e*_ + *τ*_*m*_) is shorter in *axl2*Δ*rax1*Δ and even shorter in *rsr1*Δ compared to wildtype. Hence, the timing of the initiation of polarization with respect to bud emergence is affected in both mutants. The slopes of the mean traces suggest that Cdc42 polarization occurs faster in both spatial cue mutants. Indeed, there is a significant difference in signal increase rate *r* for both mutants compared to wildtype (Fig. 3(d)). It is known that the Cdc42 polarization time in spatial cue mutants tends to be shorter, since the first phase of early Cdc42 activation by the landmark proteins is skipped [10, 34]. However, the fact that the effective rate of localization increases in both cases is surprising.

At the peak stage, both mutants reach on average similar spot volumes *V* and peak intensities (not shown), indicating a similar requirement on total Cdc42 amount. Remarkably however, the mean signal density *ρ* at the peak is lower in both spatial cue mutants compared to wildtype (Fig. 3(e)), which can occur when e.g. correlation between or variance within volume and total intensity distributions differ between backgrounds. Potential explanations include sparser packing of proteins at the polar zone or a dilution of Cdc42 by other polarizing components with an increased presence at the polar zone, either quantitatively (i.e. more of the same protein species) or qualitatively (i.e. addition of different protein species).

Finally, it has been suggested that Cdc42 has an increased collective mobility in spatial cue mutants [34, 45]. Indeed, we find that spot mobility during establishment Δ*s*_*e*_ is significantly higher in both mutants compared to wildtype, indicating a larger diffusive freedom in the absence of a bias towards the bud scar perimeter during symmetry breaking (Fig. 3(f)). Moreover, we find a slightly elevated mobility Δ*s*_*m*_ in the maintenance stage of *axl2*Δ*rax1*Δ cells. Nevertheless, similarly to wildtype, the mobility is still reduced with respect to the establishment stage in all backgrounds.

Interestingly, we find differences not only between spatial cue mutants and wildtype but also between the two spatial cue mutant variants. The dynamics of *axl2*Δ*rax1*Δ cells differ in the earlier arrival time of Cdc42 with respect to bud emergence and a slightly slower polarization rate, making this mutant an intermediate between wildtype and *rsr1*Δ in some respects. The fact that both mutants do not behave identically is perhaps not surprising, since each relies on a different mechanism behind the removal of the spatial cue. Hence, although the macroscopic phenotype of random polarization is similar between the two mutants, there is still variation at the level of Cdc42 dynamics.

### C. Large perturbation: crippling positive feedback

Next, we asked to what extent our description of Cdc42 polarization is affected in the case of a larger perturbation to polarization. To this end, we chose to disrupt near-essential scaffold protein Bem1, which binds both Cdc42-GTP effector Cla4 as well as Cdc24, a guanine exchange factor (GEF) for Cdc42 that switches it to its GTP-bound active state [46–48]. Therefore, local accumulation of Cdc42-GTP can induce a Bem1-mediated positive feedback loop that leads to amplified localization and symmetry breaking [15, 39, 49], as illustrated in Fig. 4(a). Full *bem1*Δ knockouts have a greatly reduced growth rate and display morphological defects related to polarization failure, such as large unbudded cells and elongated cell shapes [27, 50, 51].

**FIG. 4.**
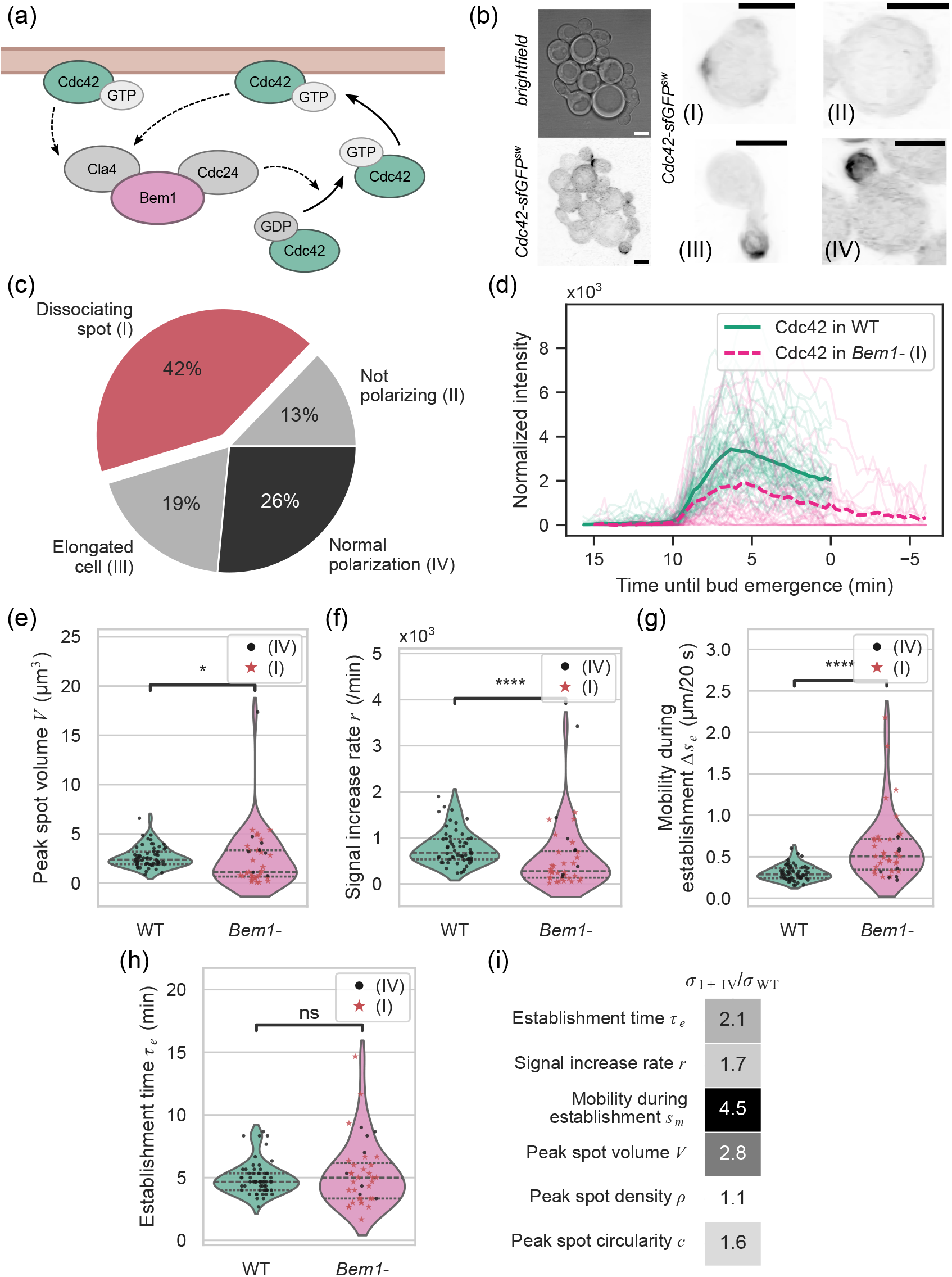
Characteristics of Cdc42 polarization after degradation of Bem1. (a) Cartoon illustrating the role of Bem1 in a positive feedback loop for Cdc42 recruitment. Cdc42 bound to GTP localizes to the cell membrane and interacts with effectors, one of which is kinase Cla4. Cla4 is one of the many interaction partners of scaffold Bem1, which also binds Cdc24, a GEF for Cdc42. This way, Bem1 recruits Cdc24 to the site of membrane-bound Cdc42-GTP, where Cdc24 locally induces more conversion of Cdc42-GDP to Cdc42-GTP, which will bind the membrane and continue the feedback loop. (b) Representative brightfield (top left) and maximum z-projected fluorescence (other panels) images of cells expressing Bem1-AID after >30 min incubation with 0.25 mM NAA (*Bem1-*). Cells exhibit defects in morphology as well as localization of Cdc42-sfGFP^sw^. Right panels show representative cells of each phenotypic category: (I) dissociating spots, (II) not polarizing, (III) polarization and growth in elongated cells and (IV) normal polarization and bud growth. Scale bars 5 µm. (c) Distribution of all observed events among the phenotypic categories illustrated in (n = 117). (d) Individual normalized intensity traces of polarized Cdc42-sfGFP^sw^ (thin lines) and mean intensity for wildtype (n = 61) and analyzable *Bem1-* cells displaying phenotype (I) (n = 31), aligned at the mean onset of polarization. (e-h) Distributions of peak spot volume *V*, signal increase rate *r*, spot mobility during establishment Δ*s*_*e*_ and establishment time *τ*_*e*_ respectively, for wildtype (n = 61) and analyzable *Bem1-* cells displaying phenotypes (I) and (IV) (n = 39). (i) Standard deviations *σ* of all relevant observable distributions for *Bem1-* cells displaying phenotypes and (IV), normalized to the respective standard deviations of the wildtype distributions. All plots: \**p* < 0.05; \*\**p* < 0.01; \*\*\**p* < 0.001; \*\*\*\**p* < 0.0001; ns, not significant (Mann-Whitney U-test). Dashed lines indicate data quartiles.

To circumvent the high likelihood of suppressor mutations in full *BEM1* knockout cells, we opted for induced degradation of Bem1 using the auxin inducible degron (AID) system [52, 53]. After endogenous labeling of Bem1 with the small AID tag, degradation can be induced by adding naphthalene acetic acid (NAA), a synthetic auxin substitute that is specifically suitable for imaging since it does not degrade into toxic products under blue light [53]. Growth rate measurements confirmed great sensitivity of the Bem1-AID strain to an NAA concentration of 0.25 mM (see Supplemental Material [29]). In the following, we incubated Bem1-AID expressing cells with 0.25 mM NAA for >30 minutes before imaging, which we refer to as Bem1-depleted cells.

As expected, polarization of Cdc42 is severely affected in cells after degradation of Bem1. Cells show significant defects in morphology and often arrest in a large, unbudded state. Inspecting the localization behavior of Cdc42-sfGFP^sw^ (Fig. 4(b)) reveals four main categories of polarization phenotypes: (I) polar spots that do not result in daughter cell growth, but instead dissociate back into the cytoplasm, (II) large cells that show no polarization during the experiment, (III) continued polarization resulting in elongated daughter cells, and (IV) seemingly normal polarization resulting in successful daughter cell growth. The largest fraction of observed events belonged to category (I), whereas complete polarization failure was relatively rare (Fig. 4(c)). This indicates that cells are generally able to break symmetry in the absence of Bem1, but fail to fully develop or maintain the spot into bud emergence.

Focusing on the dynamics of Cdc42 in dissociating spots (category (I)), the mean intensity trace shows similar general characteristics of growth and decay as in wildtype cells, but at reduced overall intensity (Fig. 4(d)). The total amount of localized Cdc42 at the peak is on average lower than in wildtype, and the intensity gradually decays down to zero rather than ending in bud emergence. This is confirmed by the smaller peak spot volume *V* as shown in Fig. 4(e), which also includes the few analyzable successful polarization events leading to bud emergence (category (IV)). While the sample size is not sufficient for strong statistical conclusions, successful polarization events seem to occur more often at larger spot volumes, although many spots with larger volumes do not lead to bud emergence. Similarly, we observe a reduction in the signal increase rate *r* in Bem1-depleted cells (Fig. 4(f)), suggesting a decreased efficiency of Cdc42 localization.

During establishment, the mobility of the spot Δ*s*_*e*_ is greatly increased in Bem1-depleted cells (Fig. 4(g)), similarly to the spatial cue mutants in Fig. 3(f). It should be noted that Bem1-depleted cells display a similar random polarization pattern as spatial cue mutants [51], meaning that polarization is not bound to the spatial landmark complex. This may explain the increase in initial mobility in both Bem1-depleted and spatial cue knockout cells. Remarkably, when considering the establishment time *τ*_*e*_, the mean timescale at which peak polarization is reached is similar in wildtype and Bem1-depleted cells (Fig. 4(h)). In other words, although the volume and signal level at the peak differ, Cdc42 accumulates on average within the same time frame in both backgrounds. This could indicate an effect of inherent timescales of negative feedback mechanisms that downregulate polarization at later stages in wildtype cells [54], which might fully destabilize the immature spots in Bem1-depleted cells.

In summary, depletion of Bem1 causes prominent disruptions to the phenotype of Cdc42 polarization, as expected. While defects arise in spot assembly and morphology, the mean time scale at which the spot is established appears to be conserved. However, this statement only concerns the population average, while the spread of the distribution in Fig. 4(h) in Bem1-depleted cells is larger than in the wildtype distribution. This observation holds true for most observables: Fig. 4(i) shows the relative magnitude of the standard deviation for each relevant observable distribution compared to the standard deviation of the corresponding wildtype distribution. Most characteristics indeed have a substantially increased variance with respect to wildtype, indicating a larger variety of phenotypes. Hence, depletion of Bem1 seems to release some constraints on Cdc42 localization and expand the spectrum of possible polarization patterns, which are most often not successful in inducing bud emergence and growth.

### D. Dynamics of other polarizing components

In the previous, we have focused on the localization of Cdc42 as the main regulator of polarity. However, we next wanted to ask whether the applicability of our descriptive framework can be extended to other polarizing protein species. For this purpose, we chose the p21 binding domain (PBD) of Gic2, a biosensor that binds Cdc42-GTP, and Spa2, a polarisome component. Gic2 by itself is an interactor of Cdc42 that plays a role in polarity establishment and septin ring assembly [55, 56]. The isolated Gic2 PBD domain binds specifically to Cdc42 in its GTP-bound state and has been extensively characterized as a biosensor for active Cdc42 [57, 58]. Hence, we expect localization of Gic2PBD to represent a subpopulation of Cdc42. Spa2 on the other hand is a member of the polarisome, a protein composite that regulates targeted exocytosis for polarized membrane deposition [20, 59]. While Spa2 is not essential for viability, knockout mutants show abnormalities in morphology and budding patterns [35, 60]. Since Spa2 is an effector of Cdc42, we opted to employ it as a case study for localization of downstream polarity components.

In the following, we use Gic2PBD-3xGFP endogenously expressed under the Gic2 promoter, integrated at a neutral chromosomal locus, as well as endogenously expressed Spa2-Citrine under its native promoter. Both constructs are integrated in wildtype backgrounds. Due to their different roles and interactions with polarization, localization of Gic2PBD and Spa2 differs slightly from Cdc42 throughout the cell cycle (Fig. 5(a)). Cdc42 localizes to the polar zone and the membrane of the daughter cell, as well as to the mother bud-neck during cytokinesis. However, while polarization is dominated by GTP-bound Cdc42, cytokinesis involves Cdc42 in its GDP-bound form [61]. Therefore, since Gic2PBD targets Cdc42-GTP, Gic2PBD follows a similar pattern of localization as Cdc42 except for cytokinesis; during this time, no localized Gic2PBD is observed. Spa2 also localizes both to the polar zone as well as the mother-bud neck during cytokinesis. Moreover, unlike for Cdc42, a localized signal can be observed in the daughter cell even after cytokinesis, which persists for some time until it dissolves prior to the next polarization event.

**FIG. 5.**
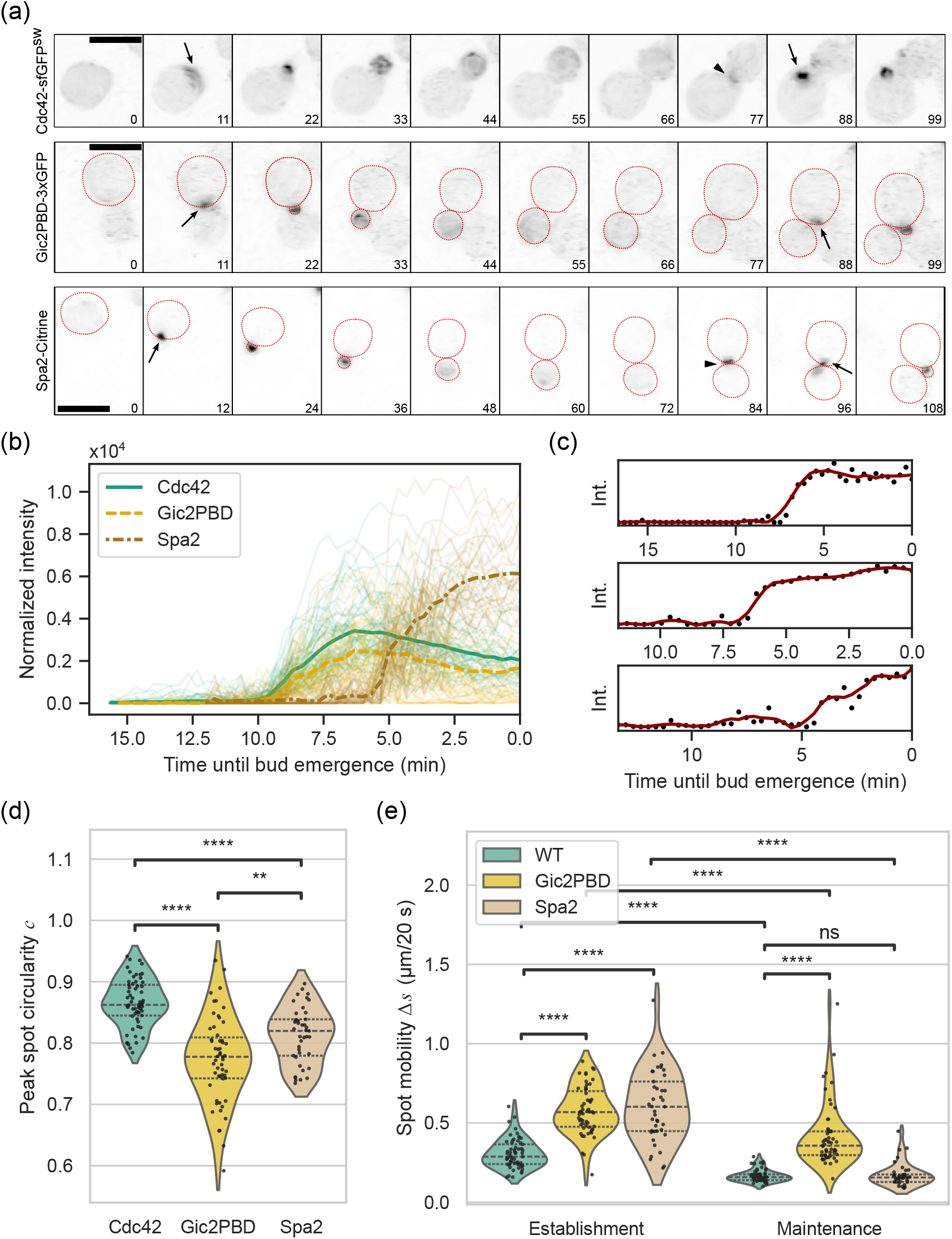
Characteristics of polarization of Gic2PBD and Spa2 in wildtype. (a) Representative time lapse max z-projected fluorescence images of Cdc42-sfGFP^sw^ (top row), Gic2PBD-3xGFP (middle row) and Spa2-Citrine (bottom row) during a full cell cycle. Arrows indicate polarization events, and arrowheads indicate localization during cytokinesis, which is absent for Gic2PBD. Numbers indicate time in minutes relative to the start of the time lapse. Cell outlines have been drawn on the images of Gic2PBD and Spa2 for clarity. Scale bars 5 µm. (b) Individual normalized intensity traces (thin lines) and mean intensity of polarized Cdc42-sfGFP^sw^ (n = 61), Gic2PBD-3xGFP (n = 55) and Spa2-Citrine (n = 41) in wildtype cells, aligned at the end of the time series corresponding to bud emergence. (c) Representative individual intensity traces of polarized Spa2-Citrine, showing polarization and stabilization (top), polarization and continued increase (middle) and step-wise polarization (bottom). (d) Distributions of peak spot circularity *c* for Cdc42, Gic2PBD and Spa2 in wildtype cells. (e) Distributions of spot mobility during establishment Δ*s*_*e*_ and during maintenance Δ*s*_*m*_ for Cdc42, Gic2PBD and Spa2 in wildtype cells. All plots: \**p* < 0.05; \*\**p* < 0.01; \*\*\**p* < 0.001; \*\*\*\**p* < 0.0001; ns, not significant (Mann-Whitney U-test). Dashed lines indicate data quartiles.

We first consider the mean intensity traces of each protein, shown in Fig. 5(b) aligned at bud emergence at the end of each trace. The dynamics of Gic2PBD are clearly strongly correlated with polarized Cdc42, in accordance with our expectations. The onset of polarization with respect to bud emergence as well as the time to reach the peak are highly similar, although the signal itself is lower on average. Spa2 on the other hand displays very different dynamics. In the mean intensity trace, onset of Spa2 polarization occurs around or after the peak of Cdc42 and Gic2PBD localization, indicating that it starts to localize at a later stage in the polarization process. This is consistent with the fact that Spa2 is considered a downstream effector of Cdc42. Moreover, the dynamics show qualitatively different behavior: rather than accumulating until a signal peak and subsequently decreasing, Spa2 localization tends to happen according to an initial rapid increase and subsequent plateau or gradually continued increase. This becomes more apparent when inspecting individual intensity traces. Figure 5(c) shows representative examples of typical behavior: initial rapid localization is either followed by a plateau or slight decrease (top) or a gradual continuation of signal increase (middle). Moreover, in certain cases polarization occurs in a more step-wise manner, showing multiple phases of increase and stabilization (bottom). For the remainder of the analysis, Spa2 polarization is considered along the same phases of establishment, peak and maintenance as Cdc42, where we define the peak as the point where the final plateau is reached; this point is identified by the exact same algorithm as before, described in the Supplemental Material [29].

Examining the morphology of the peak spot, spot volumes only differ marginally between Cdc42 and Spa2 (not shown). However, both Spa2 and particularly Gic2PBD form more irregularly shaped spots, judging by the decrease in circularity *c* (Fig. 5(d)). This is somewhat remarkable and may partially be explained by the comparatively lower signal intensity for Gic2PBD, resulting in a larger contribution of noise. However, it could also be a legitimate effect of Gic2PBD only binding to a subset of the total Cdc42 population, caused by either an inhomogeneous distribution of Cdc42-GTP and Cdc42-GDP at the polar zone or cooperative binding effects of Gic2PBD itself. The difference in circularity between Cdc42 and Spa2 is also statistically significant but smaller in absolute value. Hence, spots formed by Spa2 tend to be slightly less circular than Cdc42.

Moreover, we observe significant differences in the mobility of the spot. Gic2PBD is more mobile both during establishment and during maintenance, which may be related to its more irregular shape causing the center of mass of the signal to undergo larger displacement. Spa2 also shows greatly increased mobility during the establishment phase, but stabilizes after reaching its peak or plateau. This could be attributed to the observation that Spa2 sometimes forms multiple subclusters in close proximity to each other that alternate in strength before a single cluster is formed.

In conclusion, the quantitative properties used to describe Cdc42 polarization can also be applied to other polarizing protein species, although care needs to be taken to assess differences in dynamic behavior. Gic2PBD mostly follows the dynamics of Cdc42 as expected, although we observe differences in spot morphology and overall mobility. Spa2 localizes with different dynamic patterns, but the description along the three stages can still be applied. In general, the framework provided here is applicable to any polarizing component for which three stages of initial rapid increase, peak or plateau and subsequent spot maintenance can be distinguished. In case of qualitatively different dynamics, adaptations may have to be made to ensure that all observables remain relevant.

## III. DISCUSSION

In this work, we developed a quantitative phenomenological description of budding yeast polarization using 3D live cell microscopy, based on the dynamics of Cdc42. We derived distributions for observables characterizing each polarization stage and thus revealed quantitative bounds for polarization of Cdc42 in a wildtype background, although the distributions showed significant variability. We then applied the description to a variety of mutants, demonstrating the ability to distinguish mesoscale phenotypes between genetic backgrounds. Finally, we extended the analysis to other polarizing protein species, revealing distinct qualitative and quantitative differences in dynamics. Together, this demonstrates that our systematic framework to quantify mesoscale phenotypic properties can generate meaningful observations and aid in the formulation of novel hypotheses.

The results have several implications for the field of budding yeast polarity. So far, we have highlighted the most striking characteristics for each mutant. Putting together all observables and genetic backgrounds, we can compare the complete set of properties and identify general patterns. Figure 6 shows an overview of observable distribution means for each protein and background, normalized to the distribution means of Cdc42 polarization in wildtype. Although our list of mutants is by no means exhaustive and does not represent the full range of potential phenotypic variation, it is possible to identify some trends.

**FIG. 6.**
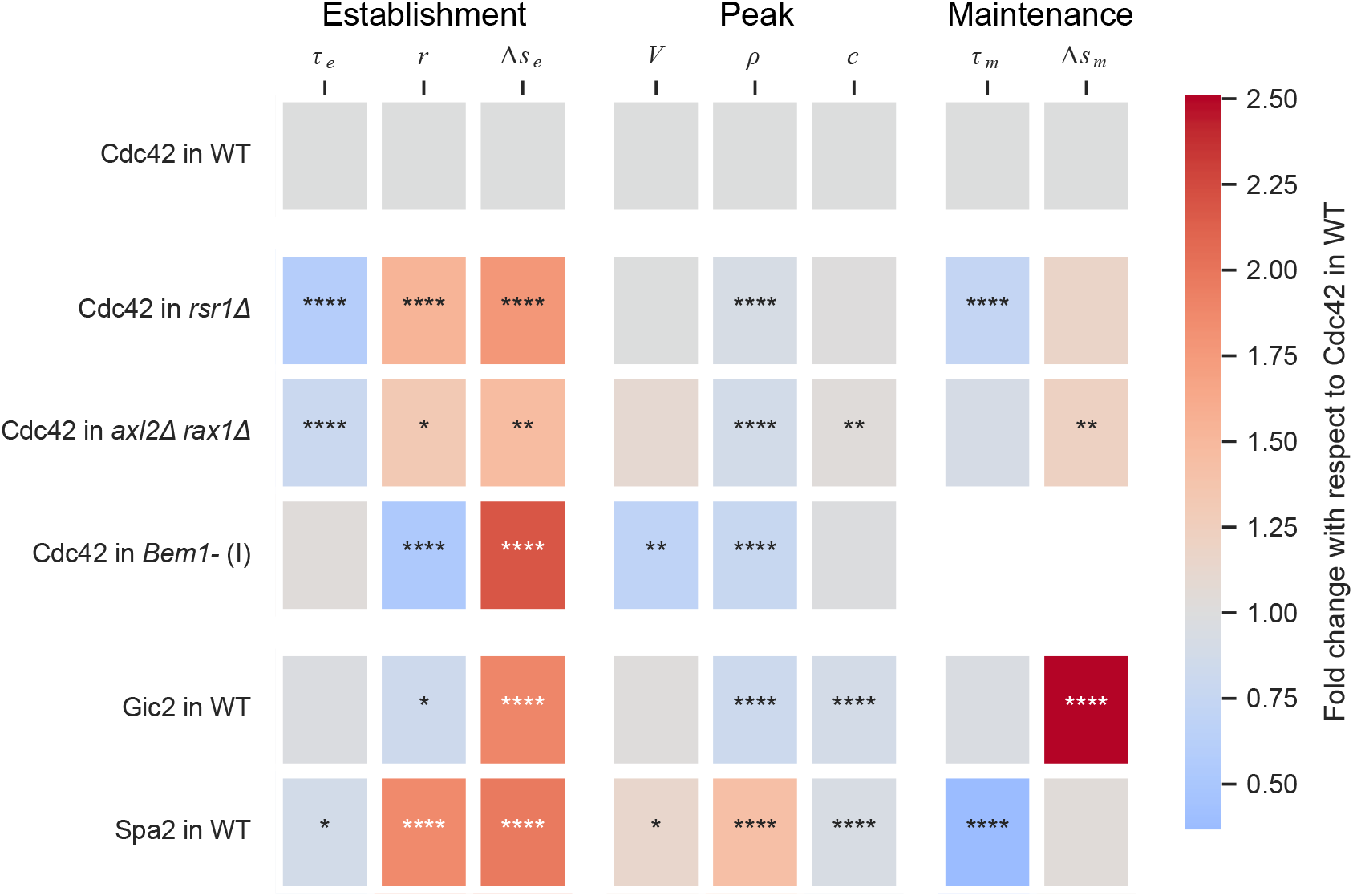
Variability of all observables in all measured backgrounds. Heatmap indicates distribution means of observables from Table I, divided by the mean of the corresponding distribution for Cdc42 in WT (reference, top row), of Cdc42 in mutant backgrounds (middle block) and other polarizing proteins in WT (bottom block). Stars indicate significance of the difference compared to the reference. \**p* < 0.05; \*\**p* < 0.01; \*\*\**p* < 0.001; \*\*\*\**p* < 0.0001; ns, not significant (Mann-Whitney U-test).

Firstly, the characteristics of polarization during the establishment stage are particularly variable. Specifically the mobility of the spot during this stage is consistently higher in all our tested backgrounds, suggesting that very precise localization of Cdc42 is not necessary for successful polarization. Stabilization of the polarity spot in asymmetric cell division is thought to occur through septin assembly at the spot periphery [58, 62], which happens at a later stage of polarization. Similarly, polarization during chemotaxis (which involves different regulatory mechanisms but many of the same components, among which is Cdc42) requires a level of mobility or wandering capacity in the polarized spot, which is driven by actin-mediated vesicle delivery and confined by septins [63–65]. Hence, we would expect relatively more diffusive freedom during the initial stages of polarization. Indeed, spot mobility decreases in the maintenance phase for each protein and mutant, although some remain more mobile than Cdc42 in wildtype. Notably, we observed that mobility in *axl2*Δ*rax1*Δ mutants remains elevated during maintenance compared to wildtype; Axl2 is known to be directly involved in septin recruitment [66]. Together, our data seem to suggest that the positioning of the polar spot is inherently mobile, but restricted by factors such as the landmark complex during establishment or septin assembly during maintenance.

Furthermore, it is interesting to note that both spatial cue mutants have a shorter establishment window. This has been previously attributed to the absence of an early Bud3-dependent Cdc42 activation phase in these mutants, leading to an overall shorter time span of polarization [34]. However, we found that the spatial cue mutants also have a higher Cdc42 localization rate. Hence, it seems that although cells in principle have access to a faster and more efficient mode of Cdc42 polarization, this is not favored in wildtype. This could be related to the fact that at least *rsr1*Δ mutants tend to have a shorter replicative and chronological lifespan [67, 68], indicating that perturbing spatial cues can come at an overall fitness cost. The reduced Cdc42 density in both spatial cue mutants could suggest a change in composition of the final spot. Spot composition can be probed by for instance proximity labeling, a technique by which proteins in the vicinity of a chosen bait protein are modified such that they can subsequently be identified through mass spectrometry [69]. This could give more insight into the origins of the faster polarization modes.

The most strongly conserved feature among the studied mutants, also considering the statistical significance of the differences between distributions, is the volume of the final spot. Notably, Bem1-depleted cells form significantly smaller spots. The fact that these smaller spots fail to produce a bud suggests that there may be a volume requirement for polarized Cdc42 to successfully initiate daughter cell emergence. Cdc42 cluster size has previously been associated with mother cell volume [70], and the spatial cue mutants studied here have indeed similar cell sizes as wildtype. It would be informative to apply our quantitative analysis to mutants with increased cell volumes (such as Cdc42 GAP mutants) to assess the effect of cell size on the other properties of polarization.

Quantification of cellular processes through imaging is regularly done for cell biological experiments, but the extent has so far been limited in the field of cell polarity. Polarization is often quantified at the cell level using measures of the magnitude of symmetry breaking [6] or in terms of polarity-specific characteristics (such as cap width or duration) that are derived situationally depending on the biological question at hand [11, 44]. A recent more generic approach was developed to be applied to polarization during shmoo formation, where a large collection of GFP-tagged proteins was systematically imaged and subjected to principal component analysis and hierarchical clustering, revealing levels of relatedness in localization patterns [71]. The variety and accessibility of image quantification methods and tools has been rapidly expanding, making systematic characterization of imaging data increasingly available for a wider scientific audience. With our method, we want to highlight the richness of information that can be deduced from relatively simple microscopy and analysis techniques.

Our phenotypic quantification of polarization can offer guidelines for future modeling efforts in several ways. The distributions of properties can be directly used to constrain modeling or simulation outcomes, by providing a physiological range for the time- and length scales that are relevant in a specific context. Moreover, qualitative aspects of polarization can serve as a point of departure for model comparison or novel theorizing. For instance, current models seem to disagree on the way initial symmetry breaking progresses into the final spot, where in some cases a broad cap is formed first that subsequently collapses and reduces in size [11] while in other cases the spot continuously expands outward from a small initial nucleation [17]. While our current data seem to favor the latter, the pattern of initial rapid establishment and subsequent maintenance and slow decay has not been reproduced by models so far and invites further investigation. Finally, the topic of biological variance is often overlooked in theoretical modeling. While some models include stochasticity in either their initial conditions or progression, they are generally unable to reproduce the level of heterogeneity in dynamics that appears in biological data. Moreover, we found that variance increased even further in the strongly perturbed Bem1-depleted mutant, raising questions about the sources of variation and their relation to the changing conditions in the mutant.

The framework and results presented here can be improved upon in several ways. Firstly, the main throughput bottleneck is currently the manual annotation of polarization events, the yield of which also strongly depends on the quality of the segmentation mask. Automating the process of selecting and checking polarized cells, either with deterministic algorithms or using machine learning, would greatly improve the throughput. It can be argued that cell synchronization protocols can also increase the yield per experiment, since this increases the number of polarizing cells within a certain time frame [72]. However, synchronization methods such as incubation with mating factor or hydroxyurea directly interfere with the cell cycle and may affect overall protein concentrations, so care should be taken in their interpretation. Apart from throughput limitations, we have so far only considered events where polarization resulted in the formation of a single spot. Competition between spots has been the subject of many studies in the past [14, 73]; it would be interesting to extend the analysis to differentiate between spatially segregated clusters. Moreover, standard image analysis such as presented here only yields relatively elementary properties, like volumes and timescales. More sophisticated techniques could be used to extract more detailed phenotypic information, such as using super-resolution techniques to assess substructure [74] or membrane wetting properties [75, 76] or using specific probes to study local crowdedness [77] or pH [78].

Polarization in budding yeast is an incredibly complex phenomenon with a deceptively simple macroscopic phenotype: asymmetric localization of cellular components. Molecular studies have contributed greatly to our understanding of the richness of interactions and mechanisms that underlie this process in lab strains. However, they also highlight the immense variability of microscopic details and the strong dependence on genetic background of these details. Phenomenological descriptions at the mesoscale, such as the framework presented here, can help identify larger scale patterns and constrain potential unifying mechanisms. The genetic variants in this work are meant as a proof of principle; application to a wider distribution of mutants would be highly valuable. Moreover, we hope to encourage the inclusion of quantitative and experimentally verifiable observables in future modeling and simulation efforts, to facilitate better comparison. Together, this can advance our understanding of the general principles underlying complex cellular processes like polarization.

## IV. METHODS

### A. Strains, plasmids and growth conditions

All yeast strains, plasmids and primers are listed in the Supplemental Material [29], as well as construction details. Fluorophore and AID tagged proteins were all integrated into their native genomic locus replacing the untagged ORF, except Gic2PBD-3xGFP, which was integrated at the HO locus under the Gic2 promoter. Yeast were grown in rich medium (YPD: 10 g/L yeast extract, Sigma-Aldrich; 20 g/L peptone, Sigma-Aldrich; 20 g/L dextrose, Sigma-Aldrich) or low fluorescence synthetic complete medium (SC-LF: 6.9 g/L nitrogen base without amino acids, folic acid and riboflavin, Formedium; 790 mg/L complete amino acid mix, Formedium; 20 g/L dextrose, Sigma-Aldrich) at 30 °C in glass culture flasks placed in a rotating incubator.

### B. Microscopy

Cells were grown to mid-log phase (OD_600_ 0.2-0.8) in SC-LF. 1 ml culture was taken out into a sterile 1.5 ml Eppendorf tube. In the case of AID degradation, cells were incubated with 0.25 mM 1-naphthaleneacetic acid (Sigma-Aldrich) at 30 °C for at least 30 minutes before continuing. Cells were spun down for 5 minutes at 3000g and 950-990 µl supernatant was removed depending on the size of the pellet. Cells were resuspended and vortexed for 1 minute before mounting 1 µl of suspension on a 35mm imaging dish with a polymer coverslip bottom (ibidi). The droplet of cells was covered by an agar pad, prepared by using a 15 ml falcon tube to cut a circular pad out of a 9 cm diameter petri dish containing 40 ml SC-LF + 3% agarose (Carl ROTH). Cells were imaged on a laser scanning confocal microscope (Eclipse Ti inverted microscope equipped with a A1R confocal unit, Nikon Instruments) with full environmental control (Tokai Hit) kept at 30 °C and a z-drift compensation module (Nikon Perfect Focus System). Images were captured using resonant scanning mode (unidirectional) with 4x line averaging (250 ms/frame), which was found to allow for sufficient signal levels as well as maintaining acquisition speed and cell viability. Excitation was done using a 488 nm (GFP) or 514 nm (Citrine) argon laser (Dynamic Laser) and emission filter 525/50 (GFP) or 540/30 (Citrine) through a 100X/1.49NA SR Apo TIRF oil objective (Nikon); signal was detected by an A1-DUG GaAsP Multi Detector unit with gain 50. Images (9 z-slices, 0.9 µm spacing, pinhole diameter 174 µm for GFP or 183 µm for Citrine) were taken every 20 s for a total of 3 hours. Images were collected from the fluorescence channel and the transmitted light detector; fluorescence images were denoised using Nikon’s Denoise.AI [79]. Acquisition parameters are justified in the Supplemental Material [29].

### C. Image analysis

Image analysis is described in detail in the Supplemental Material [29].

Cells were segmented from the background using the YeaZ GUI [31] on the ‘brightfield’ images obtained from the transmitted detector channel. Segmentation masks were first used to perform a bleaching correction on the full image stack. Individual polarization events were manually annotated from at least 10 frames before the first appearance of polarized signal until the onset of bud emergence and cropped from the full field of view by applying the YeaZ segmentation mask. Each cell stack was normalized to the average cytoplasmic intensity during the first 5 frames of the timelapse and a threshold was calculated based on the mean and standard deviation of the signal during the first 5 frames. The threshold was applied to each frame and thresholded voxels were used for the calculation of the polarization observables.

All analysis was implemented in Python.

## ACKNOWLEDGMENTS

We thank Frank van Opstal, Ramon van der Valk, Els Sweep and Esengul Yildirim for their technical support during the project. We thank Hay-Oak Park for providing the plasmid containing the Gic2PBD biosensor. We thank Marko Kaksonen for sharing his expertise on live cell imaging of budding yeast. We thank the other members of the Laan lab for the helpful discussions and feedback on the manuscript. L.L. gratefully acknowledges funding from the European Research Council under the European Union’s Horizon 2020 research and innovation programme (grant agreement 758132).

## Notes

### Competing Interest Statement

The authors have declared no competing interest.

