## Supplemental Information for "Quantitative Characterization of Budding Yeast Polarization at the Mesoscale"

October 9, 2025

### 1 Yeast strain construction

#### 1.1 Strains used in this study

| Strain name | Main text name | Relevant genotype |
|---|---|---|
| yLL3a | - | - |
| yMG01b | Cdc42-sfGFP$^{sw}$ in wildtype | *cdc42::CDC42-sfGFP*$^{sw}$*_URA3* |
| yEG01a | Cdc42-sfGFP$^{sw}$ in *rsr1*$\Delta$ | *cdc42::CDC42-sfGFP*$^{sw}$*_URA3* \\ *rsr1*$\Delta$ |
| yEG03e | Cdc42-sfGFP$^{sw}$ in *axl2*$\Delta$ *rax1*$\Delta$ | *cdc42::CDC42-sfGFP*$^{sw}$*_URA3* \\ *axl2*$\Delta$ \\ *rax1*$\Delta$ |
| yMG05a | Cdc42-sfGFP$^{sw}$ in *Bem1-* | *cdc42::CDC42-sfGFP*$^{sw}$*_URA3* \\ *bem1::BEM1-AID* \\ *YPRCtau3::OsTIR1-3xMyc* |
| yMG24c | Gic2PBD-3xGFP in wildtype | *HO::GIC2PBD-3yeGFP_URA3* |
| yLL132a | Spa2-Citrine in wildtype | *can1::Pmfa-HIS3* \\ *spa2::SPA2-Citrine_URA3* |

Table SI: Yeast strains used in this study. All strains are haploid (mating type $\alpha$ expect yLL132a, which is mating type $a$) in a W303 background. All strains have *BUD4* replaced by *BUD4* from S288c and *leu2-3*, *his3-11* and *ura3*$\Delta$.

#### 1.2 Strain construction procedures

Strain yMG01b was created by integrating *CDC42-sfGFP*$^{sw}$ at the genomic locus using homologous recombination (upstream flank 490 bp, downstream flank 310 bp) and selection by a *URA3* auxotrophic marker. A 400 bp region downstream of the ORF was left between the *CDC42* ORF and the *URA3* marker to maintain the native promoter. The *CDC42-sfGFP*$^{sw}$ sandwich construct was created by inserting superfolder GFP (sfGFP) between L134 and R384 of Cdc42, preceded by an SGGSA linker and followed by a CSGPPG linker [1].

Strain yEG01a was created from yMG01b by deletion of *RSR1* using the Cas12 gEL DNA method [2] and a 120 bp deletion fragment (60 bp upstream, 60 bp downstream).

Strain yEG03e was created from yMG01b by deletion of *AXL2* and *RAX1* simultaneously using Cas12 and 120 bp deletion fragments (60 bp upstream, 60 bp downstream).

Strain yMG05a was created from yMG01b in two steps. First, *TIR1* from *Oryza sativa* (*OsTIR1*) fused to 3xMyc with a 685 bp ADH1 promoter and 347 bp ADH1 terminator was inserted into the *YPRCtau3* locus (upstream flank 60 bp, downstream flank 60 bp) using Cas12, fully replacing *YPRCtau3*. Next, *BEM1* fused to the AID tag with a RLGGGGS linker was integrated replacing native *BEM1* (upstream flank 493 bp, downstream flank 61 bp) using Cas12, with silent PAM site mutation A491 GCT $\rightarrow$ GCG.

Strain yMG24c was created from yLL3a by integrating the *GIC2* PBD domain (*GIC2* aa1-208 with mutation W23A [3, 4]) under the native *GIC2* promoter (304 bp) fused to 3 copies of yeast optimized GFP (yeGFP) with a LINGTKAGGS linker between Gic2PBD and the first yeGFP and RS linkers between each copy of yeGFP at the *HO* locus using homologous recombination (upstream flank 909 bp, downstream flank 504 bp) and selection by a *URA3* marker. The plasmid containing Gic2PBD$^{\mathrm{W23A}}$ was kindly gifted by the Hay-Oak Park lab.

Strain yLL132a contains Spa2-Citrine with a GDGAGLIN linker integrated at the native locus using homologous recombination and selection with a *URA3* marker and was created as described in [5].

All integrations and deletions were confirmed by PCR. Inserted constructs were sequenced to exclude mutations.

#### 1.3 Validation of Bem1-AID degradation

Since absence of Bem1 leads to strong growth disruptions, we measured growth rates to verify the depletion of Bem1-AID. Cells were taken directly from glycerol stocks and suspended in 50 µl YPD. 1 µl suspension was added to 200 µl YPD in wells of a 96 well plate. Growth was monitored by measuring optical density (OD) at 600 nm using a Biotek Epoch 2 Microplate Spectrophotometer kept at 30 ℃ while shaking the plate. OD was measured every 7 minutes for >70 hours. For each strain, NAA was added to a final concentration of 0.25 mM. 6 replicates divided over 2 plates were performed for each strain and condition (with and without NAA). The resulting growth curves are shown in Fig. S1(a).

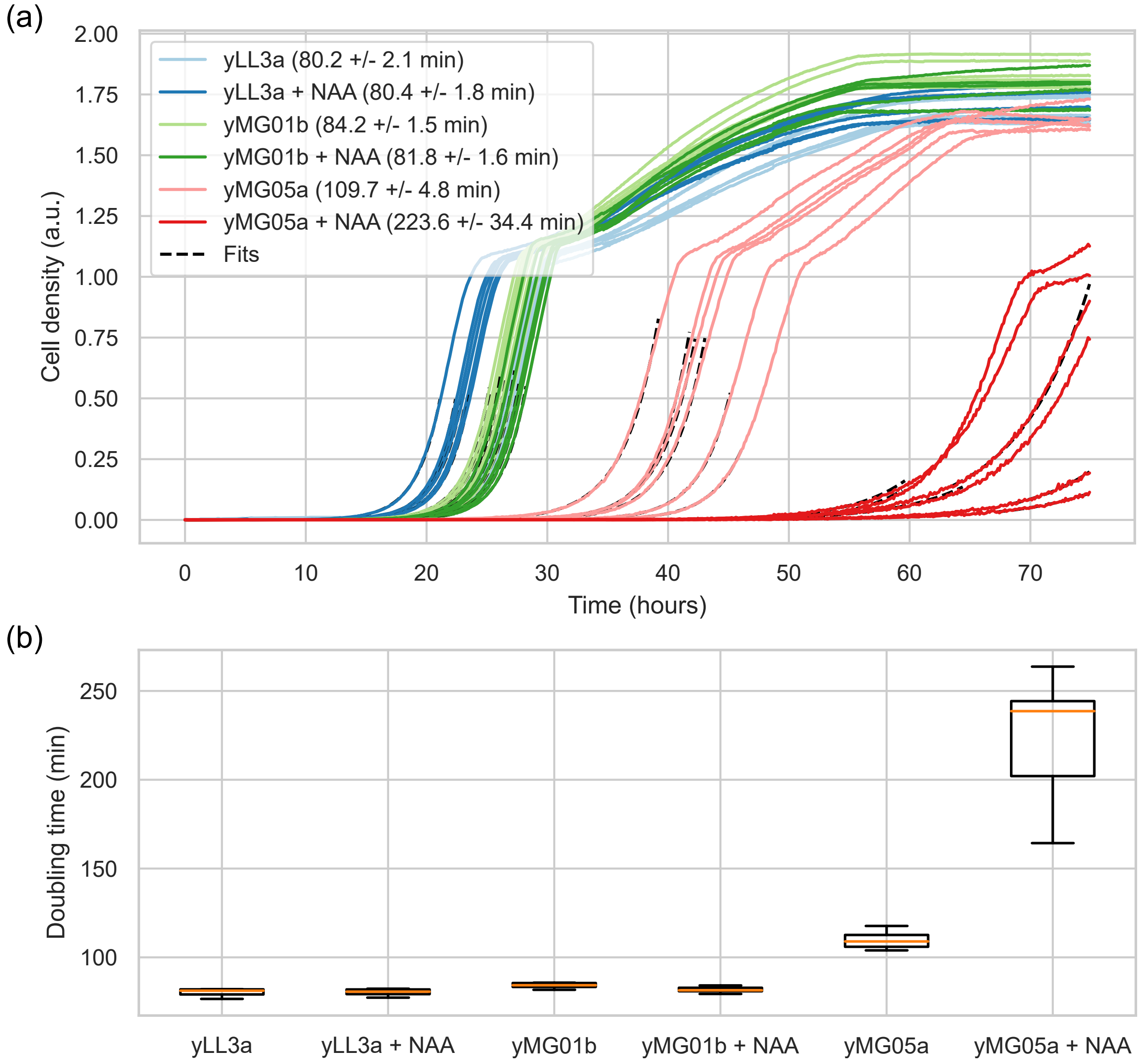

FIG. S1: Validation of Bem1-depletion phenotype using the AID system. (a) Growth curves of relevant strains: yLL3a (WT), yMG01b (Cdc42-sfGFP$^{sw}$ in WT) and yMG05a ((Cdc42-sfGFP$^{sw}$ and Bem1-AID), with and without 0.25 mM NAA. (b) Strain doubling times extracted from exponential fits to the log phase of the growth curves in (a).

To determine the doubling time during the exponential growth phase, the log of the data was fitted to a linear relation of the form $y = ax + b$ in a sliding window. The doubling time $T$ was derived from the maximum slope $a$ as $T = \log(2)/a$. Fits are shown in Fig. S1(a) as black dotted lines. Doubling times for all tested strains and conditions are shown in Fig. S1(b). From these data, we conclude that this concentration of NAA does not affect growth of wildtype cells (yLL3a) nor cells expressing Cdc42-sfGFP$^{sw}$ (yMG01b), while growth of cells expressing Cdc42-sfGFP$^{sw}$ and Bem1-AID (yMG05a) is severely impaired, providing strong evidence of Bem1 depletion. Growth of yMG05a without NAA is also slightly impaired; we hypothesize that this is caused by leaky degradation of AID-tagged protein in absence of NAA [6]. However, since the effect is small, we do not expect significant risk of suppressors in this strain.

### 2 Imaging details

All code used in this work will be available in a Github repository.

#### 2.1 Image preprocessing

Cells are segmented from the background using the YeaZ GUI [7], available from `https://github.com/rahi-lab/YeaZ-GUI`. The segmentation masks, which are obtained from brightfield images, are then applied to the fluorescence imaging data. An example of a brightfield field of view used for segmentation is shown in Fig. S2(a), along with the mask applied to the corresponding fluorescence image. Segmentation masks are first used to perform a bleaching correction on the full image stack by fitting the average maximum z-projected fluorescence signal inside the cell mask to an exponential decay of the form $I(t) = c_1 + c_2 e^{-t/c_3}$ with $c_{1-3}$ fitting parameters. (Fig. S2(b)). The raw stack is then normalized to the fitted exponential.

Individual polarization events are manually annotated by recording the mask identifier of the cell and the time window in which the event occurs, from at least 10 frames before the first appearance of polarized signal until the onset of bud emergence. Polarizing cells are then cropped from the full field of view by applying the segmentation mask to each slice of the z-stack in the annotated time window. Since the segmentation mask is 2D, this yields cylindrically shaped cell stacks (Fig. S2(c)). The cell z-stacks are then interpolated in the z-direction such that voxels have equal dimensions in each spatial direction, to account for the lower sampling rate in the axial compared to the planar direction. To compensate for uneven illumination and/or variance in laser intensity, cell stacks are normalized with respect to the average cytoplasmic intensity in the unpolarized state, calculated by taking the mean signal intensity of the maximum z-projection of the first 5 frames of the timelapse.

#### 2.2 Threshold rationalization

To identify the signal belonging to the polar spot, a threshold is calculated for each individual cell stack. Since we reasoned that the intensity of the polar spot is a significant departure from normal intensity fluctuations in the unpolarized state, we chose to base the threshold on the mean $\mu$ and standard deviation $\sigma$ of the unpolarized cytoplasmic signal, calculated by considering the max z-projected signal in the first 5 frames of the timelapse. The threshold is then defined as a number of standard deviations $k$ above the mean unpolarized intensity: $\theta = \mu + k\sigma$. To determine an appropriate value for $k$, we considered the intensity histograms of cells in unpolarized versus polarized state. An example of a representative cell is shown in Fig. S3(a), where thresholds for various values of $k$ are plotted as vertical lines on the histogram; the corresponding contours are drawn on the fluorescence image on the right. Note that while the example cell is shown in 2D (i.e. a maximum z-projection), in reality the threshold is applied in 3D. We also applied the same thresholds to a copy of the image stack that was Gaussian smoothed ($\sigma_{x,y,z,t} = (1, 1, 1, 0)$ pixels) and used the obtained contours on the unsmoothed image, which is shown in Fig. S3(b). With smoothing applied, the histogram shows better distinction between the polarized and unpolarized state and the projected contours have fewer noisy inclusions.

To quantitatively assess the effect of different threshold levels, we calculated the fraction of voxels passing the threshold as a function of $k$, both in the unpolarized state (Fig. S3(a)) and in the polarized

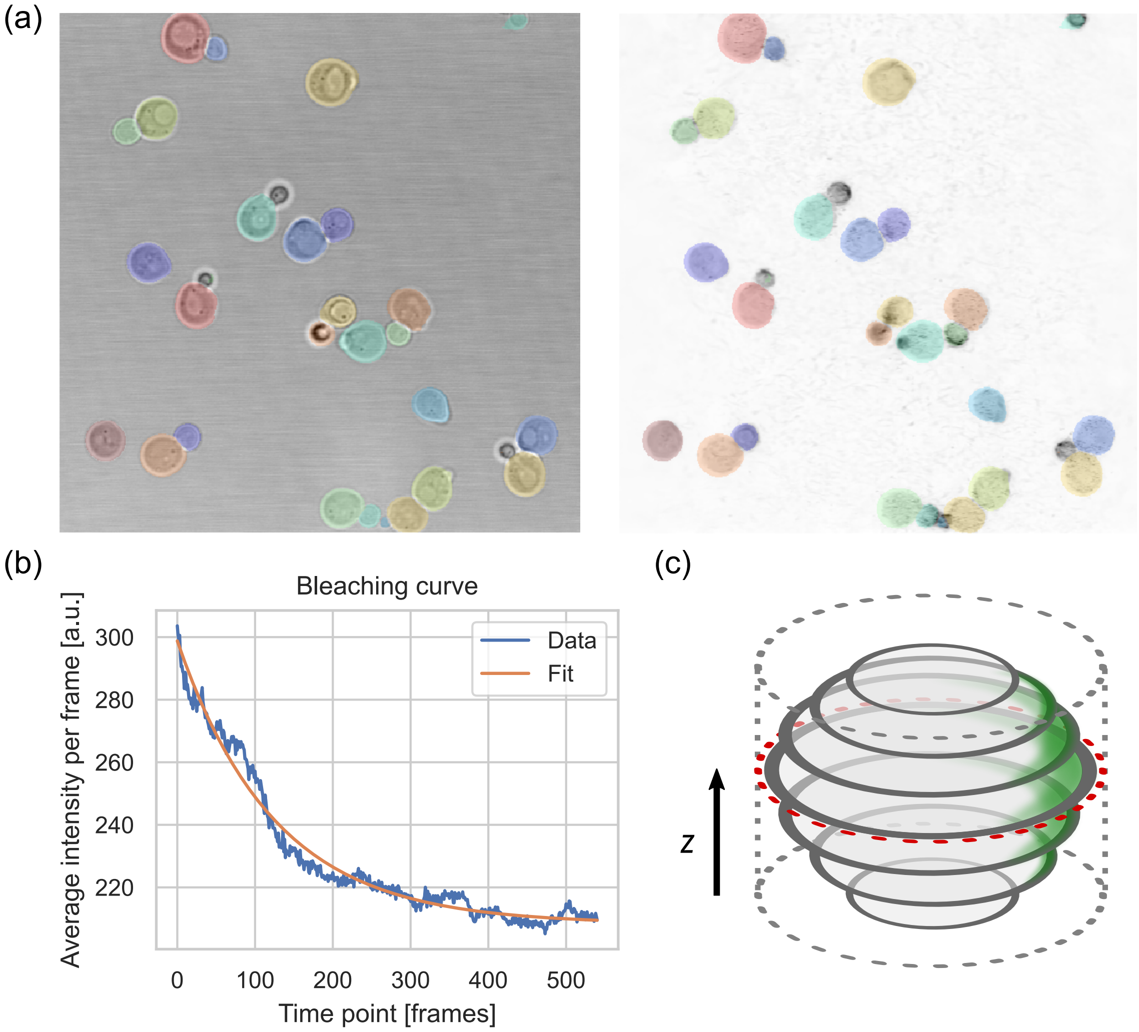

FIG. S2: Image preprocessing steps. (a) Example brightfield (left) and corresponding maximum projected fluorescence (right) image frames with the YeaZ segmentation mask applied. (b) Intensity curve used for bleaching correction of a representational experiment. (c) Illustration of the application of the 2D segmentation mask to a 3D cell stack, resulting in a cylindrical crop.

state (Fig. S3(b)), averaged over all polarization events of Cdc42-sfGFP$^{\mathrm{sw}}$ in wildtype (n = 61). We did this both for masks obtained by thresholding the raw data (unsmoothed) as well as thresholding a Gaussian smoothed copy of the data. In the unpolarized state, the desired included voxel fraction is 0% (since the cell is not polarized, no voxels should be assigned to the polar spot). As Fig. S3(c) shows, this point is reached at a lower threshold value when smoothing is applied than without smoothing. With smoothed masking, the fraction of unwanted inclusions is effectively reduced to zero at $k = 3$. In the polarized state, the difference between unsmoothed and smoothed masking is only minor and mostly independent of $k$. To maximize signal retention in the polarized state, we choose the smallest value of $k$ for which the thresholded fraction in the unpolarized state is close to zero, which occurs at $k = 3$ using smoothed masking.

Hence, for each cell, the threshold is defined as $\theta = \mu + 3\sigma$. The threshold is used to obtain a mask to identify the polarized voxels by applying it to a smoothed copy of the data. It is important to note that the smoothed copy is only used for obtaining the polarity mask; all subsequent calculations are performed on the unsmoothed data.

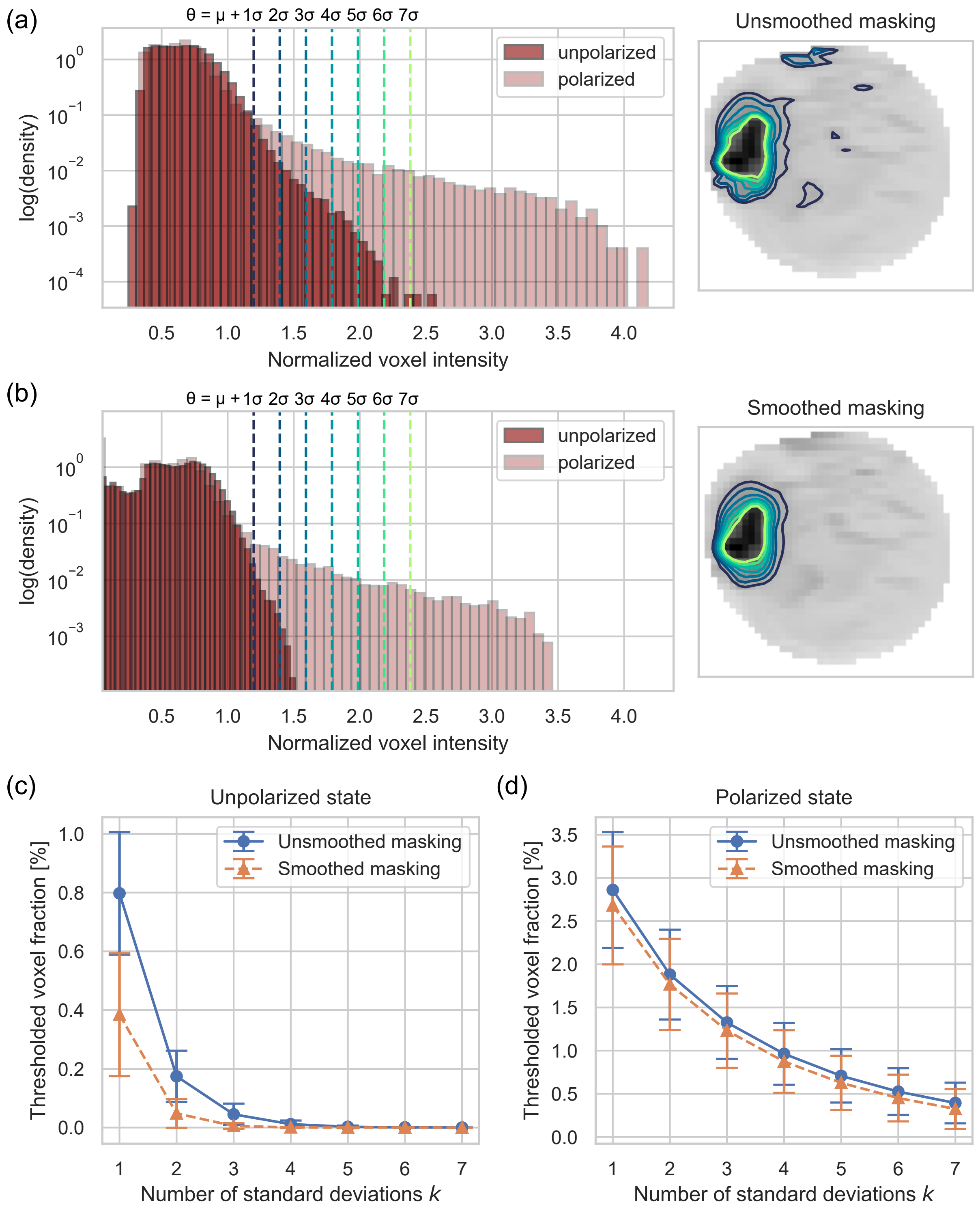

FIG. S3: Determining the threshold for identification of polarized signal. (a) Left: histogram of voxel intensities of a representative cell in unpolarized and polarized state. Thresholds are calculated as $\theta = \mu + k\sigma$, with $\mu$ and $\sigma$ the mean and standard deviation of the signal in the unpolarized state respectively. Threshold values for different $k$ are plotted on the histogram as vertical lines. Right: maximum z-projected fluorescence image of the same cell. Contour lines correspond to the signal segmented by the respective threshold values from the histogram.

FIG. S3: (continued) (b) Same as (a), but the same thresholds are now applied to the stack after Gaussian smoothing with $\sigma_{x,y,z} = 1$. (c) Average percentage of thresholded voxels of all Cdc42-sfGFP$^{sw}$ in wildtype events (n = 61) in the unpolarized state as a function of the number of standard deviations $k$ in the threshold value, for masking without and with Gaussian smoothing applied. (d) Same as (c), but for cells in the polarized state.

#### 2.3 Calculation of observables

All observables are calculated based on 3D images. For the total intensity traces, the signal inside the thresholded mask is summed for each frame in the timelapse. Smoothed curves are obtained by applying a Savitzky-Golay filter of order 3 with a window length of 1/5th of the total number of frames in the timelapse. The start of polarity establishment is then determined by identifying the global signal maximum and moving backward in time until the smoothed signal dropped below 10% of the maximum. To determine the end of polarity establishment, the first order derivative of the smoothed signal is taken. The gradient is scanned starting from the onset of polarization to find the first point where it drops below 1% of its maximum and the intensity signal is at least 80% of its maximum, indicating the local signal maximum reached after polarity establishment. The procedure is illustrated in Fig. S4(a).

Alignment of intensity traces from cells with the same genetic background is done based on the start of polarity establishment. Since not all traces have the same length, a meaningful window needs to be determined in which the mean intensity curve will be calculated. This window is set to run from the moment at which at least 50% of individual intensity curves are present, until the average total duration of polarization until bud emergence (i.e. the sum of $\tau_e$ and $\tau_m$) has elapsed (see Fig. S4(b)). For the comparison of arrival times between different genetic backgrounds, the mean intensity curves for each background are aligned with respect to each other by bud emergence at the end of the mean intensity profile.

Observables are calculated individually for each polarization event. The determined start and end points of polarization are used to divide the process into three stages: the establishment stage runs between the start and end point of polarization, the peak stage is defined as the five frames around the end point of polarization, and the maintenance stage runs from the end point of polarization until the end of the timelapse defined by bud emergence. The establishment time $\tau_e$ is then calculated by taking the time difference between the beginning and the end of polarity establishment. Signal increase rate $r$ is calculated by dividing the average total intensity in the peak frames by the establishment time. The spot mobility during establishment $\Delta s_e$ is calculated by taking the Euclidean distance between the signal center of mass for each consecutive frame during establishment and taking the average of all step sizes.

The peak spot volume $V$ is calculated by summing the volume of thresholded voxels for the peak frames and taking the average. The peak spot density is calculated by dividing the average total polarized intensity in the peak frames by the average peak spot volume. Peak spot circularity is calculated by rotating the center of mass vector of the polarity signal during the peak frames to align with the z-direction and taking a maximum z-projection (see Fig. S4(c)). Circularity is then defined as the average circularity of the 2D contours of the projected peak frames calculated as $c = \frac{4\pi A}{L^2}$, with $A$ the area within the contour and $L$ the length of the contour.

The spot mobility during maintenance $\Delta s_m$ is calculated in the same way as $\Delta s_e$, but for the frames belonging to the maintenance stage. The maintenance time $\tau_m$ is calculated by taking the time difference between the calculated end point of polarity establishment and the end of the annotated timelapse.

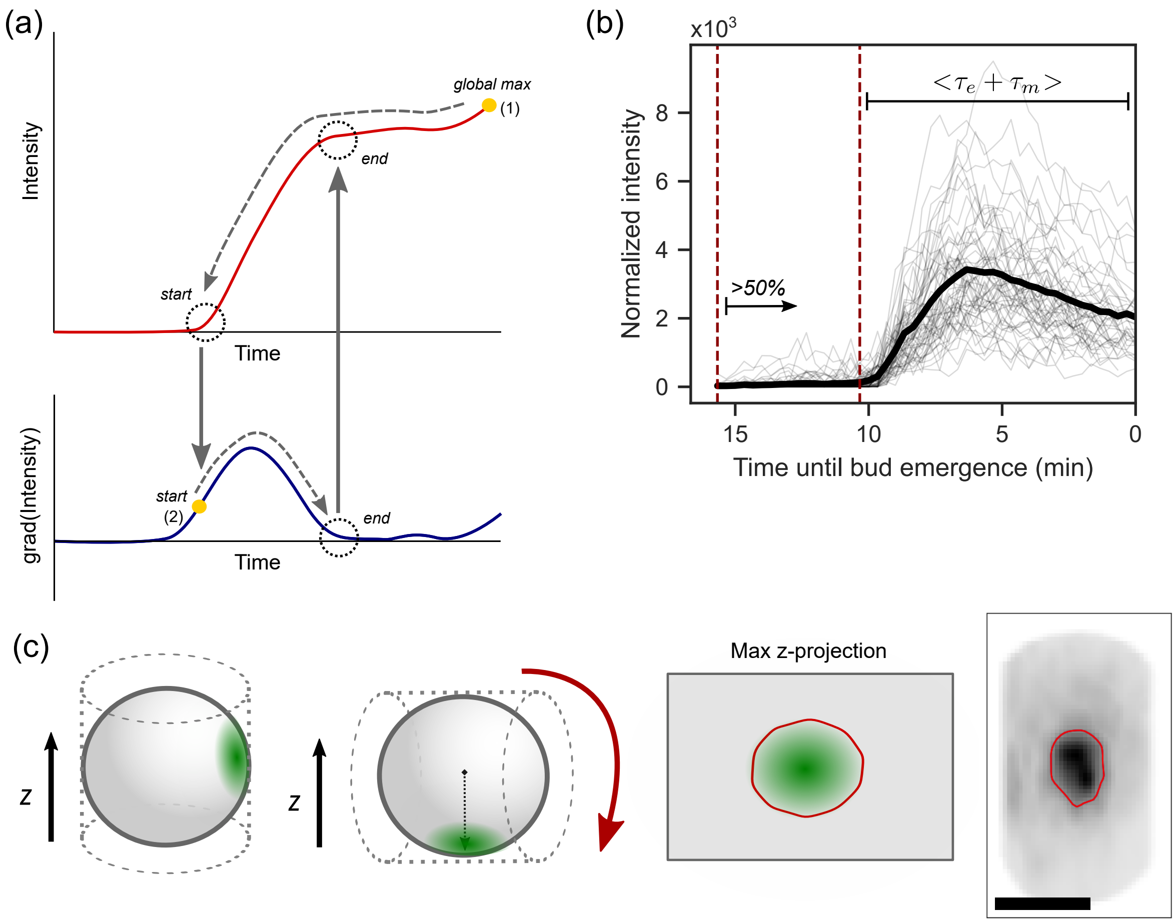

FIG. S4: Illustrations of observable calculations. (a) Schematic representations of a typical polar intensity curve (top) and its first order derivative (bottom). First, the global maximum of the signal (1) is determined and the signal is scanned backward to identify the start of polarization. Then, the derivative is scanned from the start of polarization (2) to identify the next (near) local minimum which defines end of polarization. (b) Illustration of intensity trace alignment and cut-off. Individual traces are aligned at polarity onset. The mean intensity is calculated starting at the point where $>$50% of traces are present and ending after the mean duration of polarity before bud emergence, i.e. the sum of $\tau_e$ and $\tau_m$. (c) Schematic illustration of the procedure to calculate circularity. Cell stacks are rotated such that the center of mass of the polar spot is directed towards the z-axis and a maximum projection is taken in the z-direction. Circularity is calculated from the 2D contour applied to the projected frame, using the same threshold value as used for the 3D stack. Bottom right, example of a projected image of a representative cell (scale bar 3 µm).

### 3 Imaging controls

#### 3.1 Cell viability under imaging conditions

Cells were imaged as described in the main text. Live cell imaging involves a trade-off between signal levels and cell viability. To check the viability of the cells in our imaging conditions, we estimated doubling times from the image data by counting the number of cells in the segmentation mask in each frame and fitting the cell count as a function of time to exponential growth, similar to the procedure in section 1.3. An example of a fit is shown in Fig. S5(a). We performed this analysis for our two imaging conditions: GFP, for which we analyzed 3 replicates of Cdc42-sfGFP in wildtype (strain yMG01b), and Citrine, for which we analyzed 5 replicates of Spa2-Citrine in wildtype (strain yES02a). Resulting doubling times are shown in Fig. S5(b).

We found slightly higher doubling times than those found when growing cells in YPD (see Fig. S1). This can have several explanations. Firstly, cells grown in rich medium are expected to have

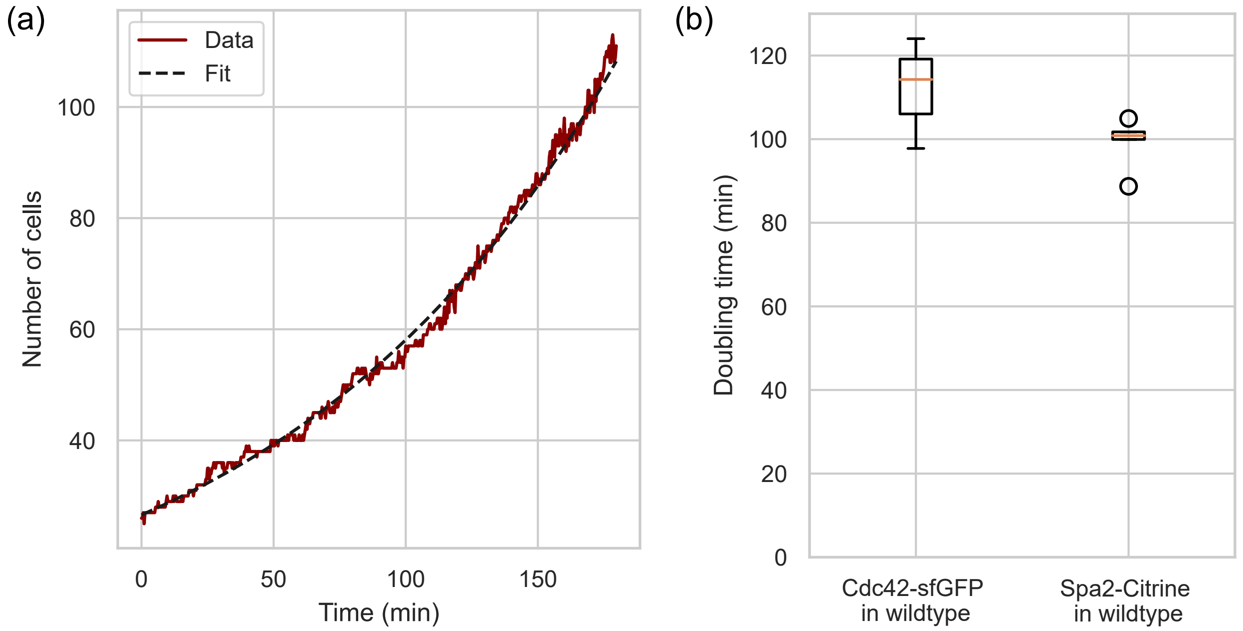

FIG. S5: Doubling times in microscopy conditions. (a) Representative example of cell count data from a microscopy experiment based on the segmentation mask (red) fitted to an exponential curve (black dashed line). (b) Doubling times extracted from microscopy data of Cdc42-sfGFP in wildtype and Spa2-Citrine in wildtype respectively.

faster growth rates than cells grown in synthetic minimal medium used for microscopy. Secondly, the algorithm to calculate doubling times from growth curve data used in section 1.3 scans the curve to find the highest growth rate, while the fits here are performed over a fixed window equal to the full duration of the experiment. Finally, yeast is particularly sensitive to exposure to blue light used for the excitation of GFP, which may also explain the slightly faster doubling times for Spa2-Citrine, since Citrine is excited at a higher wavelength. Overall however, the increases in doubling time are relatively minor and cells divide normally, hence we conclude that our imaging conditions are sufficiently permissive.

##### 3.2 Effect of image denoising

To increase the signal-to-background ratio and limit the effects of Shot noise to which resonant scanning confocal microscopy is particularly sensitive, raw image data is passed through the Denoise.ai algorithm developed by Nikon. To check for unwanted artifacts, we discuss the consequences of applying Denoise.ai to the subsequent analysis outcomes for Cdc42 in wildtype.

Figure S6(a) shows representative image frames from a cell in polarized state, projected towards the polar spot (see the calculation of circularity in section 2.3), without (left column) and with (right column) Denoise.ai. The red contour in the bottom two frames indicates the calculated threshold level (as discussed in 2.2). Comparing the images, applying Denoise.ai reduces background noise, while the overall shape of the signal is preserved. The calculated threshold compared to the maximum signal level is slightly higher, as is visible from the narrower contour in the bottom frame; with Denoise.ai, the threshold contour is wider and more signal is included. This is also evident from the intensity curves in Fig. S6(b). While the qualitative characteristics of the dynamics are conserved, the overall signal level is improved significantly by applying Denoise.ai.

A consequence of the higher signal levels after denoising is the fact that the onset of polarization can be detected at an earlier stage. This also means that the establishment time $\tau_e$ is estimated to be shorter without Denoise.ai (Fig. S6(c)), since it takes more time for the initial weak signal to exceed

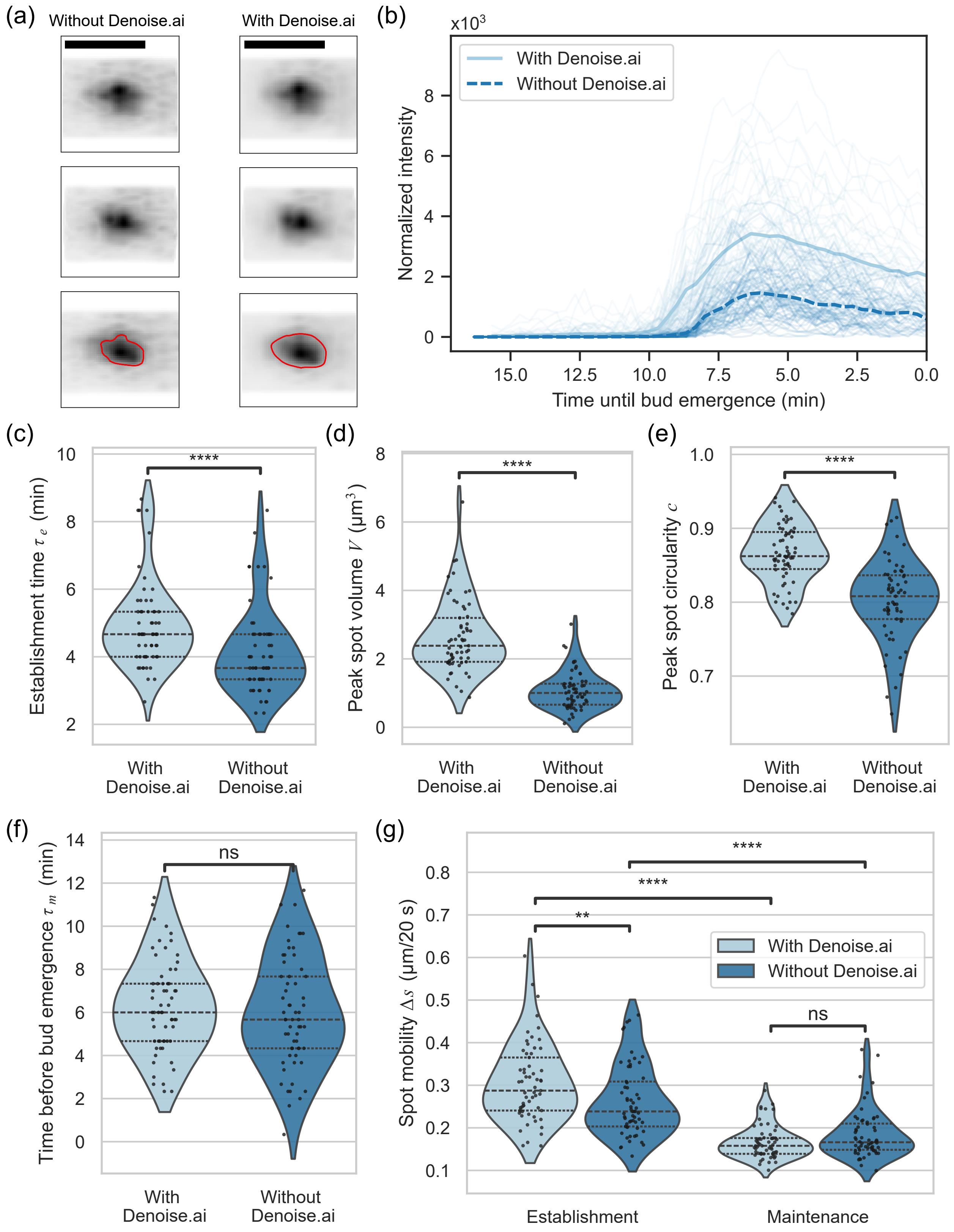

FIG. S6: Effect of Denoise.ai on the analysis of Cdc42 in wildtype. (a) Example microscopy images without (left column) and with (right column) Denoise.ai applied, rotated and projected towards the polar spot. Bottom row shows default threshold contours in red lines. Scale bars 5 μm (b) Individual normalized intensity traces of polarized Cdc42-sfGFP$^{sw}$ in wildtype (thin lines, n = 61) and mean intensity with and without Denoise.ai applied. (c-g) Distributions of establishment time $\tau_e$, volume $V$, circularity $c$, time before bud emergence $\tau_m$ and mobility during establishment/maintenance $\Delta s_{e,m}$ respectively, with and without Denoise.ai applied. All plots: *$p < 0.05$; **$p < 0.01$; ***$p < 0.001$; ****$p < 0.0001$; ns, not significant (Mann-Whitney U-test). Dashed lines indicate data quartiles.

the threshold. Similarly, the calculated total spot volume is larger when applying Denoise.ai (Fig. S6(d)). Another consequence of the wider spot contours is an increase in circularity of the final spot, since contours are less smoothed out at lower signal-to-noise levels (Fig. S6(e)). This implies that applying Denoise.ai might make it more difficult to find differences in circularity between populations, since more detailed features can be missed.

Several general characteristics are mostly unaffected. Figure S6 shows that the time before bud emergence $\tau_m$ remains similar after applying Denoise.ai. Moreover, spot mobility trends are conserved: although mobility during establishment $\Delta s_e$ is estimated slightly higher with Denoise.ai, mobility during maintenance $\Delta s_m$ is unaffected, and the stabilization of the spot during maintenance is observed in both cases.

Hence, while applying Denoise.ai results in some expected differences between calculated observables, key features are unaffected. Since denoising images increases the amount of signal that can be detected, which becomes more relevant for cases in which the polarity signal is weak (such as Bem1-depleted cells), we chose to apply it for all our image processing. Importantly, the analysis behind our quantitative framework is most effective for making comparisons between phenotypes, rather than producing absolute numbers. When denoising does not lead to obvious artifacts and is consistently applied to all data, which is the case here, valid comparisons can be made.
